# Draft genome of six Cuban *Anolis* lizards and insights into genetic changes during the diversification

**DOI:** 10.1101/2022.05.06.490966

**Authors:** Shunsuke Kanamori, Luis M. Díaz, Antonio Cádiz, Katsushi Yamaguchi, Shuji Shigenobu, Masakado Kawata

## Abstract

The detection of various type of genomic variants and their accumulation processes during species diversification and adaptive radiation is important for understanding the molecular and genetic basis of evolution. *Anolis* lizards in the West Indies are good models for studying the mechanism of the evolution because of the repeated evolution of their morphology and the ecology. In this study, we performed *de novo* genome assembly of six Cuban *Anolis* lizards with different ecomorphs and thermal habitats (*Anolis isolepis, Anolis allisoni, Anolis porcatus, Anolis allogus, Anolis homolechis*, and *Anolis sagrei*). As a result, we obtained six novel draft genomes with relatively long and high gene completeness, with scaffold N50 ranging from 5.56–39.79 Mb, and vertebrate Benchmarking Universal Single-Copy Orthologs completeness ranging from 77.5% to 86.9%. Subsequently, we performed comparative analysis of genomic contents including those of mainland *Anolis* lizards to estimate genetic variations that had emerged and accumulated during the diversification of *Anolis* lizards. Comparing the repeat element compositions and repeat landscapes revealed differences in the accumulation process between Cuban trunk-crown and trunk-ground species, LTR accumulation observed only in *A. carolinensis*, and separate expansions of several families of LINE in each of Cuban trunk-ground species. The analysis of duplicated genes suggested that the proportional difference of duplicated gene number among Cuban *Anolis* lizards may be associated to the difference of their habitat range. Furthermore, Pairwise Sequentially Markovian Coalescent analysis proposed that the effective population sizes of each species might have been affected by Cuba’s geohistory. Hence, these six novel draft genome assemblies and detected genetic variations can be a springboard for the further genetic elucidation of the *Anolis* lizard’s diversification.

**Significance:** *Anolis* lizard in the West Indies is excellent model for studying the mechanisms of speciation and adaptive evolution. Still, due to a lack of genome assemblies, genetic variations and accumulation process of them involved in the diversification remain largely unexplored. In this study, we reported the novel genome assemblies of six Cuban *Anolis* lizards and analyzed evolution of genome contents. From comparative genomic analysis and inferences of genetic variation accumulation process, we detected species- and lineage-specific transposon accumulation processes and gene copy number evolution, considered to be associated with the adaptation to their habitats. Additionally, we estimated past effective population sizes and the results suggested its relationship to Cuba’s geohistory.

## Introduction

Elucidating how genetic variation emerges and accumulates during species diversification is important for understanding the mechanisms of creating species diversity and adaptive evolution. With the recent development of sequencing technologies, genomes of various taxonomic groups other than model organisms, such as humans and mice, have been sequenced. In some lineages, the genomes of a large number of species have been sequenced and assembled, and progress has been made in elucidating details concerning the emergence and accumulation processes of mutations, leading to a better understanding adaptive evolution and diversity creation mechanisms within lineages (Brawand et al. 2014; Lamichhaney et al. 2015; Feng et al. 2020; McGee et al. 2020). Parallel evolution and adaptive radiation are important subjects in investigating the mechanisms of species diversification and adaptive evolution. Galapagos finches and African cichlids are good models that have undergone such diversifications and whose genomes, including those of many of their species, have been sequenced and analyzed. In Galapagos finches, genomic loci commonly involved in the evolution of their bill shape have been identified and it has been suggested that introducing allele through introgressive hybridization between different populations within species have also contributed to their responses to recent environmental changes (Lamichhaney et al. 2015). In African cichlids, it has been shown that ancestrally generated polymorphisms are involved in the adaptive evolution of repeated body color and morphology, and that insertions of transposable elements and gene duplications occur rapidly in the ancestors, some of which have also been detected to be associated with the evolution of gene expression (Brawand et al. 2014). Moreover, the densification of genome-assembled species within their lineages and genome-wide association analyses, have shown that indels accumulate in their ancestors and are associated with their diet and habitat (McGee et al. 2020).

*Anolis* lizards of the West Indies, as well as Galapagos finches and African cichlids, are model organisms for parallel evolution and adaptive radiation. Phylogenetically distant species with similar physical structures in their habitats often show similar morphology and behavior (reviewed in Losos 2009) among them. These adaptive evolution of morphology and ecology to physical structure of their habitats has been estimated to occur multiple times independently (e.g., Losos et al. 1998; Mahler et al. 2013). The common set of physical structure that they use, morphology, and behavior is called an “ecomorph,” such as crown-giant, trunk-crown, twig, trunk, trunk-ground, and grass-bush (reviewed in Losos 2009). Moreover, thermal habitat is also diverse and there is high diversity in the body temperature among these species (e.g., Hertz et al. 2013; Gunderson et al. 2018; reviewed in Losos 2009). Differences in thermal habitats often occur between closely related species (Ruibal 1961; Schettino 1999; Schettino et al. 2010; Cádiz et al. 2013; Gunderson et al. 2018), suggesting that adaptive evolution to different thermal habitats also occurred repeatedly. Therefore, elucidating creation and accumulation processes of genetic mutations during the parallel evolution and adaptive radiation of *Anolis* lizards in the West Indies is expected to yield important insights into the genetic or molecular basis of the morphological characteristics, physiology and ecology of lizards, their mechanism and the repeatability of evolution, and the relationship between genetic mutation, evolvability, and evolutionary constraints.

The genome assembly of *Anolis carolinensis*, the only species of the *Anolis* lizard native to the United States, has been reconstructed (Alföldi et al. 2011). In *A. carolinensis*, population analysis of their genome-wide SNPs, using this genome as a reference, has been conducted to estimate regions under natural selection by winter storms (Campbell-Staton et al. 2017), and transcriptome data has been analyzed to detect changes in gene expression related to cold tolerance (Campbell-Staton et al. 2018), suggesting that genes involved in neural activity (Campbell-Staton et al. 2017) and blood coagulation (Campbell-Staton et al. 2018) are involved in cold adaptation. For the *Anolis cybote* group in Hispaniola, genes involved in adaptation to the climate have been identified by detecting genes with climate-correlated SNPs using genome annotation of *A. carolinensis* (Rodríguez et al. 2017). For *Anolis cristatellus* in Puerto Rico, RNA-seq reads of the species were mapped to the genome of *A. carolinensis* and genome-wide SNP analysis was used to infer regions of differentiation between urban and forest populations. As a result, a non-synonymous SNP in the gene encoding arginyl-transfer RNA synthetase (RARS) was shown to be associated with heat tolerance in urban populations (Campbell-staton et al. 2020). The results of mapping sequence reads derived from the genomes of species of various ecomorph to the genome of *A. carolinensis* and examining the convergence of protein sequences show that the convergence of protein sequences does not reflect the convergence of ecomorphs (Corbett-Detig et al. 2020). In addition, the genome assemblies of three Central American *Anolis* lizards (*Anolis frenatus, Anolis auratus* and *Anolis apletophallus*) have already been reconstructed. Their comparative analyses, including the genome of *A. carolinensis*, detect positive selection in *hox* genes and genes involved in olfactory reception and reproduction (Tollis et al. 2018). Moreover, highly complete chromosome-scale genome assembly of *Anolis sagrei ordinatus*, a subspecies of *A. sagrei* collected in the Bahamas, has recently been reported (Geneva et al. 2021). Like *A. carolinensis, A. sagrei* is an excellent model among *Anolis* lizards for evolutionary and ecological studies, and further advances in the study of the genetic basis of adaptive evolution, plasticity, and their invasiveness through comparative analyses among populations are expected. Moreover, *hox* gene clusters have been sequenced in several *Anolis* lizards of mainland and the West Indies and their sequences and *hox* gene expression have been examined, suggesting that the insertion of transposable elements in the *hox* cluster affects *hox* gene expression, and is related to speciation events (Feiner 2016, 2019). However, the genome assembly of most *Anolis* lizards of the West Indies, a good model for studying convergent evolution and adaptive radiation, has not been reconstructed, and little is known about the emergence and the accumulation processes of genetic variation other than point mutations and especially genetic variations in non-coding regions during adaptive evolution.

Cuba hosts 65 species of *Anolis* lizards, the largest number of species in the West Indies islands, and their ecomorphs and thermal habitats are diverse, with species belonging to all six ecomorphs and unique ecomorph (Losos 2009; Cádiz et al. 2018). Species belonging to the same ecomorph construct monophyletic lineages (Cádiz et al. 2018). Closely related species within the monophyletic lineages have different thermal habitats with varying tree covers, air temperatures, and degrees of exposure to sunlight (Ruibal 1961; Schettino 1999; Schettino et al. 2010; Cádiz et al. 2013). For example, three trunk-ground ecomorph species are closely related: *Anolis allogus, Anolis sagrei*, and *Anolis homolechis. A. allogus* inhabits relatively cool forests and does not bask, while *A. sagrei* inhabits open areas outside the forest, where there is more sun, and frequently basks under direct sunlight, and *A. homolechis* inhabits forest margins, where temperatures are intermediate, and basks under filtered sun (Schettino 1999; Schettino et al. 2010; Cádiz et al. 2013). Besides, the degree to which they thrive in urban areas also varies by species. Some species are rarely found in urban areas, while *Anolis porcatus, Anolis allisoni* and *A. sagrei*, which naturally inhabit hot and open areas, thrive in urban areas (Schettino 1999; Schettino et al. 2010; Winchell et al. 2020). Several species of *Anolis* lizard have been reported to have been introduced and colonized in regions other than their native areas. For instance, *A. sagrei*, native to Cuba, has invaded Florida and Taiwan (Kolbe et al. 2004), and *A. carolinensis*, a close relative of *A. porcatus*, native to Cuba, has invaded Okinawa and Ogasawara Islands and Hawaii Islands from its native mainland in the USA (Suzuki-Ohno et al. 2017; Tamate et al. 2017). In *Anolis* lizards of the West Indies, species that experience hot and dry conditions in their natural habitat have been suggested to tend to be more tolerant of urban environments (Winchell et al. 2020). Therefore, elucidating the genetic variation that has occurred among Cuban *Anolis* lizards will help us to understand the adaptive evolutionary mechanisms of ecomorph, thermal habitat, urban tolerance, and invasiveness. In Cuban *Anolis* lizards, the evolutionary mechanisms of adaptation to thermal habitats have been studied at the genetic level by comparing three trunk-ground species that inhabit different thermal habitats, *A. allogus, A. homolechis*, and *A. sagrei*. Akashi et al. (2016) analyzed the transcriptome expression and observed that the expression of genes related to circadian rhythms and protein translation varied with rearing temperature. Akashi et al. (2018) also showed that the temperature-triggering escape behavior from heat and the activation of a molecular heat sensor, TRPA1 in *A. sagrei* and *A. homolechis* were higher than those in *A. allogus*. Moreover, phylogenetic analysis of several Cuban *Anolis* lizard’s coding sequences suggested that genes involved in stress response and cardiac function had undergone positive selection as *A. allisoni* and *A. porcatus* and *A. sagrei* adapted to open areas, and that these genes may be related to their current thriving in urban areas (Kanamori et al. 2021). Recently, genome editing technology has also been established in *A. sagrei* (Rasys et al. 2019). It will accelerate our understanding of the genetic basis of adaptive evolution in *Anolis* lizards. However, the genome assembly of the Cuban *Anolis* lizards has not been reconstructed, and most of the genetic variations involved in adaptive evolution have remained undetected.

In this study, we reconstructed novel draft genome assemblies of six species of Cuban *Anolis* lizard species: three closely related trunk-crown species (forest-dwelling *Anolis isolepis*, hot and open area-dwelling *A. allisoni*, and *A. porcatus*) and three closely related trunk-ground species (forest-dwelling *A. allogus*, hot and open area-dwelling *A. sagrei*, and forest margins-dwelling *A. homolechis*) and investigated their differences in genomic characteristics, such as the composition of genes and transposable elements, between the species. Although genome assembly of *A. sagrei ordinatus* from the Bahamas, a subspecies of *A. sagrei*, has already been reported (Geneva et al. 2021), *A. sagrei* has been identified to be of Cuban origin (Reynolds et al. 2020). Hence, we analyzed the genome assembly of the Cuban *A. sagrei* (*A. sagrei sagrei*) to unify the reconstruction and annotation methods of genome assemblies to be compared, and to detect genetic variations in the formation of common ecological *A. sagrei* features acquired before their introduction to other regions. In addition, the genome assemblies were used to estimate the change processes of past effective population sizes of each species. Results from the analysis of these novel draft genome assemblies will provide essential information to further elucidate the mechanism of diversification and adaptive evolution of *Anolis* lizards.

## Materials and Methods

### Sample preparation, genome sequencing, and *de novo* assembly

Specimens for all *Anolis* species were collected from Cuba: individuals for *A. isolepis, A. allogus* and *A. sagrei* were sampled from Las Terrazas, Artemisa, *A. porcatus* from Topes de Collantes, Sancti Spíritus, *A. homolechis* from Macambo, Guantánamo, and *A. allisoni* from Trinidad, Sancti Spíritus. Sample collection in Cuba, exportation to Japan the use for the research were approved by the Centro de Control y Gestión Ambiental of the Agencia de Medio Ambiente de Cuba. One individual for each species was collected. Individuals for *A. isolepis, A. porcatus*, and *A. sagrei* were females and individuals for *A. allisoni, A. allogus*, and *A. homolechis* were males. DNA was extracted from the brain for *A. isolepis, A. porcatus, A. allisoni* and *A. homolechis* and the muscle for *A. sagrei* and *A. allogus* according to HMW gDNA Extraction Protocol in the Sample Preparation Demonstrated Protocol provided by 10x Genomics®. Library preparation was performed using Chromium Controller (10x Genomics®, Pleasanton, CA, USA) according to Genome Reagent Kits v2 User Guide provided by 10x Genomics®, after which genome libraries were sequenced by Illumina HiSeq X (2 × 150-bp reads). The yielded reads for each genome were assembled *de novo* using Supernova v.2.1.1 (Weisenfeld et al. 2017), with an ideal coverage of ∼56× (Weisenfeld et al. 2017). First, we conducted the assembly with all the yielded reads. Then, if Supernova reported that the coverage was >56×, we calculated the number of reads that would bring the coverage to about 56× using the estimated genome size and conducted the assembly again, setting the number of reads to be used as such. Haplotigs and heterozygous overlapping duplications found in the assemblies were removed in the following steps. First, to obtain cleaned sequence reads, 10x barcodes in the sequence reads were removed using Longranger basic (https://support.10xgenomics.com/genome-exome/software/pipelines/latest/what-is-long-ranger), after which NGS QC Toolkit (Patel & Jain 2012) was used to perform quality cuts on the reads. The clean reads were then mapped back to each assembled genome using BWA (Li & Durbin 2009) and Minimap2 (Li 2018). Haplotigs and overlaps in the genomes were detected and removed using purge_dups (Guan et al. 2020) based on the depth of short read alignments. The gene completeness of genome assemblies before and after the removal of haplotigs and overlaps were assessed by examining the completeness of Benchmarking Universal Single-copy Orthologs (BUSCOs) for vertebrates using BUSCO software (Simão et al. 2015). As shown in the results section, the percentage of duplicated vertebrate BUSCOs decreased after the removing haplotigs and overlaps. Therefore, we decided that those with haplotigs and overlaps removed would be the final assemblies, and we performed subsequent analyses using them.

### Annotation of repeat elements and drawing repeat landscapes

For comparison of the composition of repeat elements in the genome among the species and survey of the repeat element accumulation process, we detected repeat elements in the genome assemblies and estimated historical dynamics of repeat element accumulation in the following steps. First, repeat elements in all *Anolis* genomes included in this study were searched *de novo* using RepeatModeler (Smit et al. 2008-2015). Next, the resulting library of *de novo* repeat elements was integrated with the library of the *Anolis* repeat sequences already deposited in Repbase (Bao et al. 2015). The integrated library of the repeat elements was finally used to detect repeat elements in the genomes and categorize them into classes and families using RepeatMasker (Smit et al. 2013–2015). Afterward, detected repeat elements were aligned to consensus sequences for each family by RepeatMasker and Kimura’s two-parameter distances (K2P distance) among them were calculated using calcDivergenceFromAlign.pl in the RepeatMasker package. Subsequently, the histogram based on calculated K2P values (repeat landscape) was plotted for each genome using createRepeatLandscape.pl from the RepeatMasker package to estimate the historical dynamics of the repeat element accumulation, and we converted K2P distance of repeat landscape to years using parseRM.pl (available at https://github.com/4ureliek/Parsing-RepeatMasker-Outputs, Kapusta et al. 2017). We used the average of DNA substitution rates estimated for *Anolis* lizard branches for each species repeat landscape conversion, as described in the “Estimation of DNA substitution rate and divergence time” section.

### Gene annotation, ortholog grouping, and duplicated gene identification

Gene models for repeat masked genome assemblies of *A. isolepis, A. allisoni, A. porcatus, A. allogus, A. homolechis*, and *A. sagrei* were constructed using BRAKER v2.1.6 (Brůna et al. 2021) with RNA-seq data and protein sequences of *A. carolinensis* obtained from Ensemble release 104. RNA-seq reads of each *Anolis* lizard (Akashi et al. 2016; Kanamori et al. 2021), available from the DDBJ Sequence Read Archive (DRA) of the DNA Data Bank of Japan (DDBJ) (Supplementary Table 1), were aligned to each genome assembly using GSNAP (Wu & Nacu 2010). Other nondefault options set when we run BRAKER were “--etpmode, --softmasking, --gff3.” BLAST search was used to identify predicted protein coding genes that did not match any protein sequence of UniProtKB/Swiss-Prot database, and Funannotate v1.8.7 (Palmer & Stajich 2020) was used to filter out such genes, predict UTR regions, and annotate gene names and Gene Ontology (GO) terms for each gene.

For identifying duplicated genes, predicted genes were clustered into ortholog groups (orthogroups) using OrthoFinder v.2.5.4 (Emms & Kelly 2015, 2019) using the longest protein sequences of genes predicted from the genomes of *A. isolepis, A. allisoni, A. porcatus, A. allogus, A. homolechis, A. sagrei*, and *Pogona vitticeps*, which was selected as the outgroup. The sequences of *P. vitticeps* were retrieved from Ensembl release 104. When multiple proteins from a single species were grouped into the same orthogroup, genes coding these proteins were considered duplicated genes. We then calculated the proportion of duplicated genes (*P*_D_: number of duplicated genes / total number of genes) for each species. We considered that differences in annotation methods and genome assembly quality would make comparisons of gene numbers and *P*_D_ difficult. Therefore we included only six Cuban *Anolis* lizards, for which genome assembly and annotation conducted in this study, for comparative analysis of duplicate genes. The independence of the ratio of duplicated to single gene numbers and species was tested separately for three closely related Cuban trunk-crown *Anolis* lizards (*A. isolepis, A. allisoni*, and *A. porcatus*) and three closely related Cuban trunk-ground *Anolis* lizards (*A. allogus, A. homolechis*, and *A. sagrei*) by G-test with a simultaneous test procedure (STP, Sokal and Rohlf 1994) to see whether there were significant differences in *P*_D_ among species, and if so, which species had significant differences in *P*_D_.

### Estimation of DNA substitution rate and divergence time

We used a 4-fold degenerate (4D) site of single copy ortholog genes shared by a total 41 species of sarcopterygian vertebrate species to reconstruct phylogenetic trees and estimate DNA substitution rates and divergence times of *Anolis* species. These 41 sarcopterygian vertebrates include 34 species selected from the major groups of amniotes (mammals, birds, crocodilians, turtles, and squamates), six species of Cuban *Anolis* lizards that we reconstructed their genome assembly in this study, and a coelacanth as the outgroup (Supplementary Table 2). The protein sequences of those species were obtained from databases listed in Supplementary Table 2. The coding sequences of *Gekko japonicus, A. apletophallus, A. auratus*, and *A. frenatus* were extracted from their genome assemblies referring annotation data using GffRead (Pertea & Pertea 2020). Databases from which coding sequences of other species were obtained are also listed in supplementary Table 2. A total of 41 species were analyzed, including six Cuban *Anolis* lizards for which genome assemblies were reconstructed in this study. Their single copy orthologs were grouped using SonicParanoid (Cosentino & Iwasaki 2019), comparing longest protein sequences for each gene. The coding sequences of single copy orthologs were aligned for each orthogroup with the codon model using PRANK (Löytynoja & Goldman 2005), and 4D sites shared by 80% of these species were extracted from the alignment. We estimated DNA substitution rate and divergence time by Bayesian relaxed molecular clock approach using MCMCTree in PAML package version 4.9j (Yang 2007). In advance, substitution rate were roughly estimated using baseml in PAML package version 4.8 (Yang 2007), with one fixed calibration point for the root: 418 mya (Benton et al. 2015), to set up a prior distribution of substitution rate. Then, MCMCTree was run to sample 10,000 times, with sampling frequency set to 10,000, after a burn-in of 25,000,000 iterations. Maximum likelihood (ML) tree was reconstructed using RAxML (Stamatakis 2014) with GTR+Γ model. We used this ML tree without branch length as the input tree. Calibration points were set as previously presented by Benton et al. (2015) as follows: 418 million years ago (mya) for the fixed root age; 256–296 mya for Diapsida; 247–260 mya for Archosauria; 169– 210 mya for Squamata; 165–202 mya for Mammalia; and 66–87 mya for Neognathae. The average of DNA substitution rates for *Anolis* lizard branches and each rate for these species terminal branch obtained from this analysis were used to estimate the time scale of the accumulation processes of transposable elements, and the effective population size history, respectively.

### Estimation of effective population size in the past

The past effective population size of six Cuban *Anolis* lizards analyzed in this study was estimated using the Pairwise Sequentially Markovian Coalescent (PSMC) model (Li & Durbin 2011). After the removing 10x barcodes and conducting quality cuts, sequence reads were mapped back to corresponding genome assemblies of each species using BWA (Li & Durbin 2009). SNPs were called in regions, where reads were mapped at a depth of one-third to twice the average depth for a genome, using samtools and bcftools (Li et al. 2009), following the method recommended by README of PSMC software. The input consensus sequence was generated using bcftools (Li et al. 2009) for each genome. The PSMC analyses were run with a time interval pattern and maximum 2N_0_ coalescent time set to “4 + 30 × 2 + 4 + 6 + 10” and 5, respectively, performing 100 bootstrap replicates. When rescaling the time and population size, the generation time was set to one year, and mutation rate to 1.4 × 10^−9^, 1.5 × 10^−9^, 1.8 × 10^−9^, 1.6 × 10^−9^, 2.0 × 10^−9^, and 2.1 × 10^−9^ for *A. isolepis, A. allisoni, A. porcatus, A. allogus, A. homolechis*, and *A. sagrei*, respectively, which were obtained from phylogenetic analysis.

## Results

### *De novo* genome assembly and gene annotation

Genomes of *A. isolepis, A. allisoni, A. porcatus, A. allogus, A. homolechis*, and *A. sagrei* were sequenced at 47.26–63.30× coverage (Table 1). As a result of *de novo* assembly with the number of reads adjusted so that the coverage was about 56× if it exceeds 56×, scaffold N50 and the total length of the six *de novo* genome assemblies were 5.69–42.71 Mb and 1.53–1.88 Gb, respectively (Supplementary Table 3). Genome size (Gb) estimated by Supernova v2.1.1 (Weisenfeld et al. 2017) of *A. isolepis, A. allisoni, A. porcatus, A. allogus, A. homolechis*, and *A. sagrei* were 2.08, 2.26, 2.11, 2.33, 2.66, and 2.05, respectively. The coverage of complete vertebrate BUSCOs was 79.4%–88.0%, of which duplicated vertebrate BUSCOs was 1.6%–4.3% (Supplementary Table 4). After removing the haplotigs and overlaps, scaffold N50 and total length of them were 5.56–39.79 Mb and 1.53–1.88 Gb, respectively, and the coverage of complete vertebrate BUSCOs and duplicated vertebrate BUSCOs decreased slightly to 77.5%–86.9% and 0.6%–2.2%, respectively (Table 2; Supplementary Table 4). Scaffold N50 of these genome assemblies were much shorter than chromosome level genome assembly of *A. carolinensis* (AnoCar2.0), but considerably longer than draft genome assemblies of *A. frenatus* (Afren1.0), *A. auratus* (Aaur1.0), and *A. apletophallus* (Aapl1.0) (Fig. 1). Furthermore, the coverages of BUSCOs for them was comparable to that of *A. carolinensis* genome (Anocar2.0) (Fig. 1). The GC content in the genome assemblies of *A. isolepis, A. allisoni, A. porcatus, A. allogus, A. homolechis*, and *A. sagrei* after removing the haplotigs and overlaps were 40.14%, 40.11%, 40.0%, 41.05%, 41.02%, and 40.96%, respectively, those in genome assemblies of mainland *Anolis* lizards were as follows: 40.32% for *A. carolinensis*, 49.44% for *A. frenatus*, 42.81% for *A. auratus*, and 40.92% for *A. apletophallus*. Although they were roughly equal, the GC-content of Cuban trunk-ground lineage species appeared slightly higher than that of Cuban trunk-crown lineage species and their relative, *A. carolinensis*. The difference between these lineages could also be confirmed in the frequency distribution of the GC content percentage within the 5 kb windows (Fig. 2). Additionally, genomic GC content of *A. apletophallus* is close to that of three Cuban trunk-ground species, while the distribution of GC content of *A. apletophallus* is biased toward lower GC content than those of these species. Also, in a previous study, although Tollis et al. (2018) already highlighted a higher GC bias in *A. frenatus* compared to the other three mainland species, the comparison with six Cuban species confirmed that it had the highest GC content and the distribution of GC content was also very biased toward higher GC content compared to the other species.

**Table 1.**
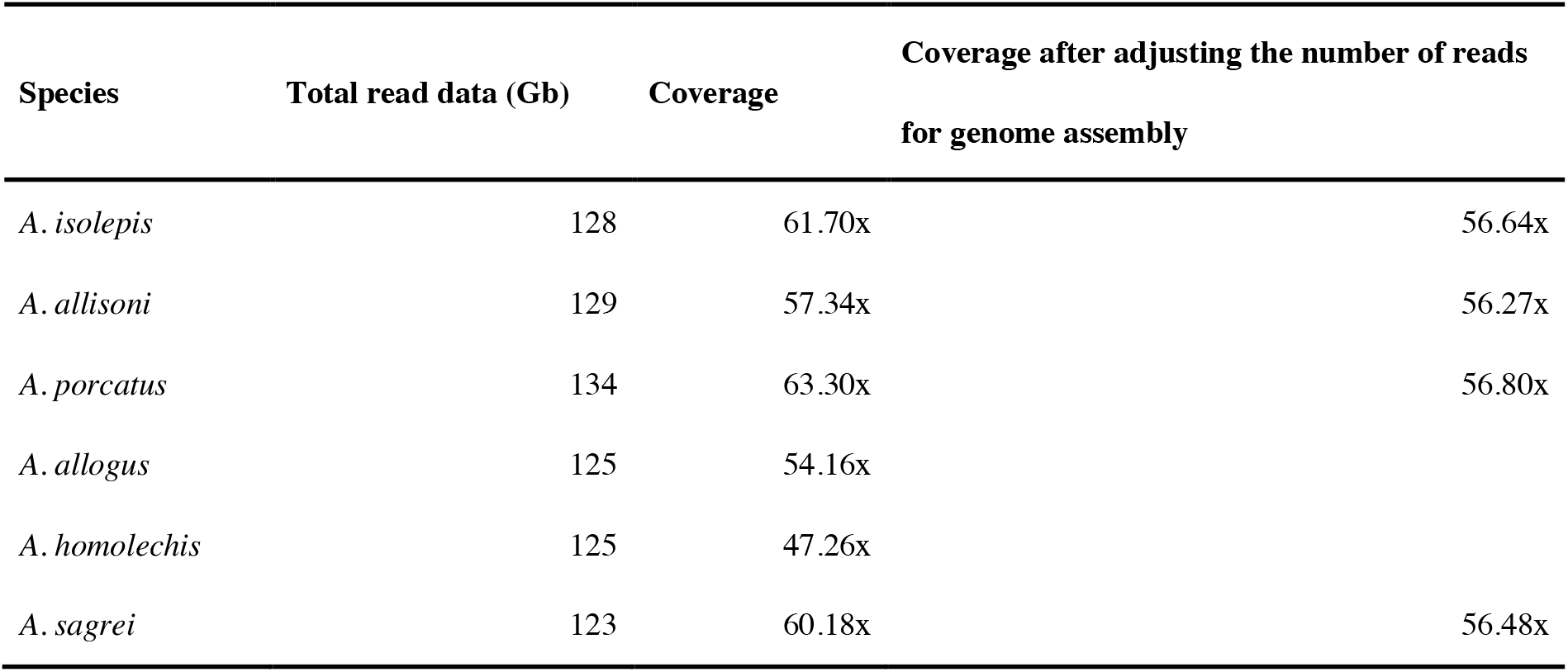
Genome sequencing results.

**Table 2.** Descriptive statistics of the reconstructed genome assemblies.

| Species | Scaffold N50 (Mb) | Total length (Gb) | Number ≥ 10Kb | BUSCO metrics | | | |
| --- | --- | --- | --- | --- | --- | --- | --- |
|  |  |  |  | Complete | (Single | Duplicated) | Fragmented |
| *A. isolepis* (Aisol1.0) | 24.70 | 1.67 | 2.24 K | 86.9% | 86.1% | 0.8% | 7.0% |
| *A. allisoni* (Aalli1.0) | 5.56 | 1.75 | 5.28 K | 83.8% | 83.1% | 0.7% | 9.4% |
| *A. porcatus* (Aporc1.0) | 23.09 | 1.74 | 5.80 K | 83.7% | 83.1% | 0.6% | 9.4% |
| *A. allogus* (Aallo1.0) | 39.79 | 2.09 | 4.68 K | 86.0% | 83.8% | 2.2% | 8.4% |
| *A. homolechis* (Ahomo1.0) | 24.32 | 1.97 | 8.60 K | 77.5% | 76.6% | 0.9% | 12.3% |
| *A. sagrei* (Asagr1.0) | 31.30 | 1.90 | 7.39 K | 85.0% | 84.0% | 1.0% | 8.9% |

**Fig. 1.**
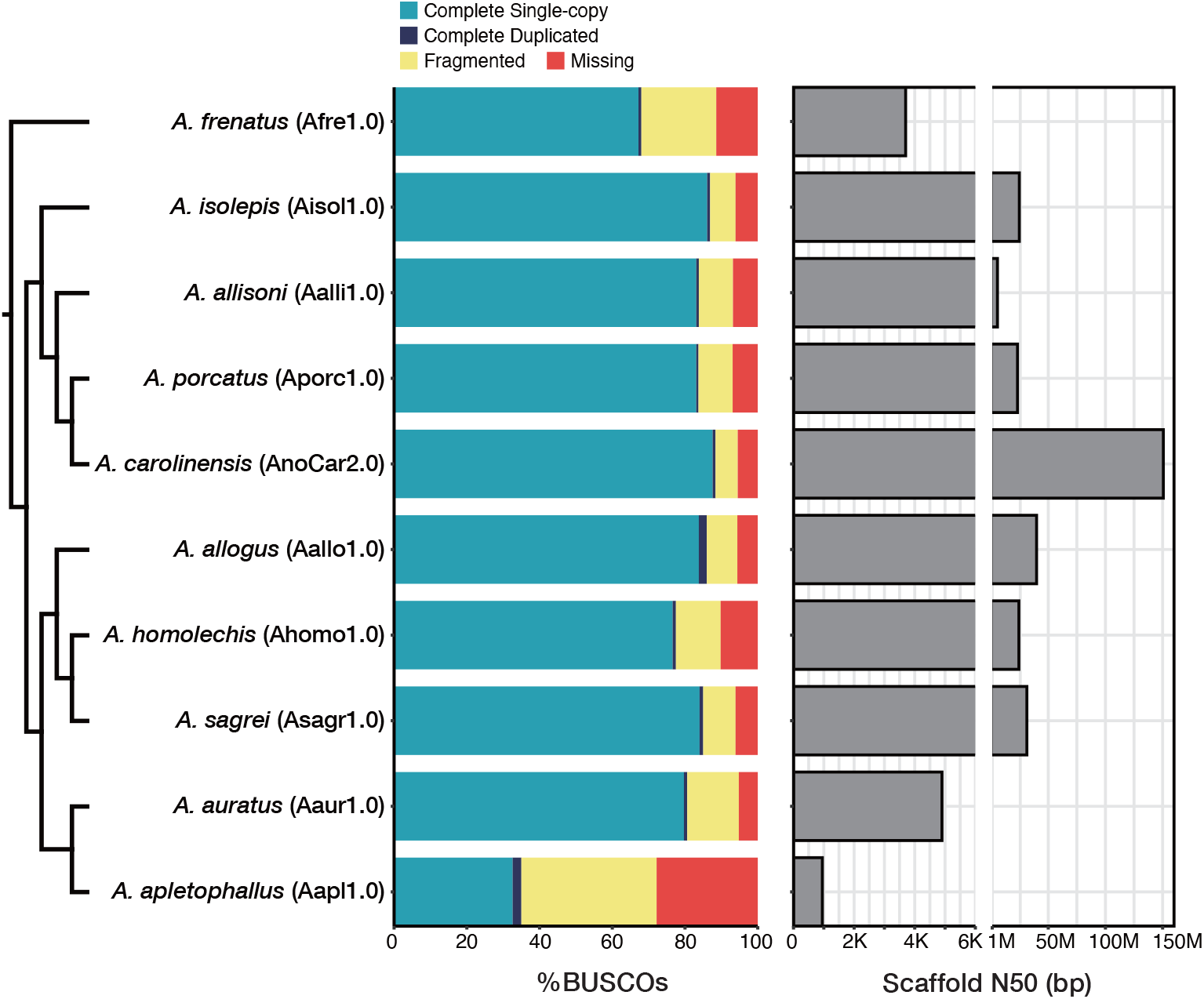
Phylogenetic relationships between *Anolis* lizards included in this study and descriptive statistics of the analyzed genome assemblies.

**Fig. 2.**
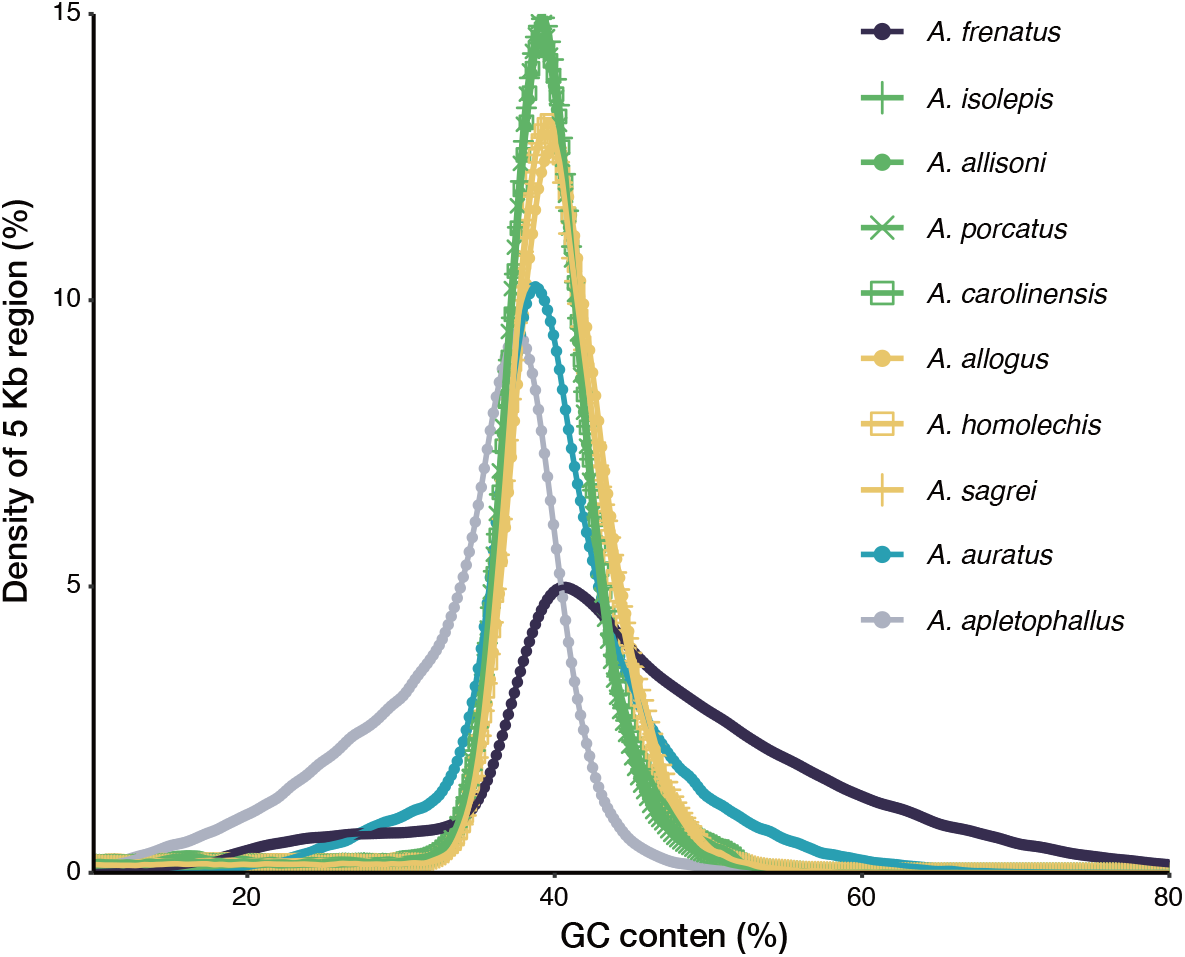
The distribution of GC content in 5 kb windows of genome assemblies for *Anolis* lizards. Green lines and points indicate the distributions for *A. isolepis, A. allisoni*, and *A. porcatus* from the Cuban trunk-crown lineage, and their relative, *A. carolinensis*; Yellow lines and points indicate the distribution for *A. allogus, A. homolechis*, and *A. sagrei* from the Cuban trunk-ground lineage; Navy blue, light blue, and gray lines and points indicate the distributions for *A. frenatus, A. auratus* and *A. apletophallus*, respectively.

Gene models were constructed on repeat masked genome assemblies after removing the haplotigs and overlaps. Then, after the filtering of genes whose encoded proteins do not match any protein in the UniProtKB/Swiss-Prot database, we predicted 21,688, 23,503, 24,100, 23,232, 25,839, and 23,819 protein coding genes for *A. isolepis, A. allisoni, A. porcatus, A. allogus, A. homolechis*, and *A. sagrei*, respectively.

### The composition of repeat elements and repeat landscape

The percentage of repeat element lengths in the genome was similar for all the six Cuban *Anolis* lizards at 37%–41% (Supplementary Tables 5–10). By the same method, we also re-annotated repeat elements for genome assemblies of three mainland *Anolis* lizards (*A. carolinensis, A. auratus*, and *A. apletophallus*), except for *A. frenatus*. Although *A. frenatus* was excluded from the analysis because of computational time constraints, we used repeat contents and repeat landscape of this species obtained from Tollis et al. (2018) analysis to compare with those of other *Anolis* lizards. The percentage of repeat element lengths in the genome of *A. carolinensis, A. auratus*, and *A. apletophallus* was also similar to Cuban *Anolis* lizards at 40%, 37%, and 27%, respectively (Supplementary Tables 11–13). That of *A. frenatus* was also similar to Cuban *Anolis* lizards at 38% (Tollis et al. 2018). Then, we compared the composition and landscape of classes and families of the repeat elements among these ten species. The results showed that genomes of the three Cuban trunk-ground species (*A. allogus, A. homolechis*, and *A. sagrei*) had more LINEs than those of other species (Fig. 3; Supplementary Tables 5–13). We also observed that the genome of *A. carolinensis* contained the most LTRs (percentage) among nine species (Fig. 3; Supplementary Tables 5–13; That for *A. frenatus* was 3.5% (Tollis et al. 2018)). Additionally, we observed a steep peak of LTR accumulation only in the repeat landscape of *A. carolinensis* (Fig. 3). According to the analysis of Tollis et al. (2018), although the *A. frenatus* genome contained more LTRs than that of *A. carolinensis* in terms of number, the steep LTR waves seen in the repeat landscape for *A. carolinensis* were lacking in that for *A. frenatus* (Tollis et al. 2018). The steep wave of LTR observed only in *A. carolinensis* was mainly composed of Gypsy family, and small waves of Pao family (Supplementary Fig. 1). Among the three Cuban trunk-ground species, which had more LINEs, different accumulation waves were observed in the repeat landscape of some LINE families (Supplementary Fig. 2). For example, *A. homolechis* and *A. sagrei* genome has more LINE-RTE-BovB than *A. allogus* genome dose, and two waves of its accumulation in *A. homolechis* and *A. sagrei* genomes were insignificant for *A. allogus* (Supplementary Fig. 2).

**Fig. 3.**
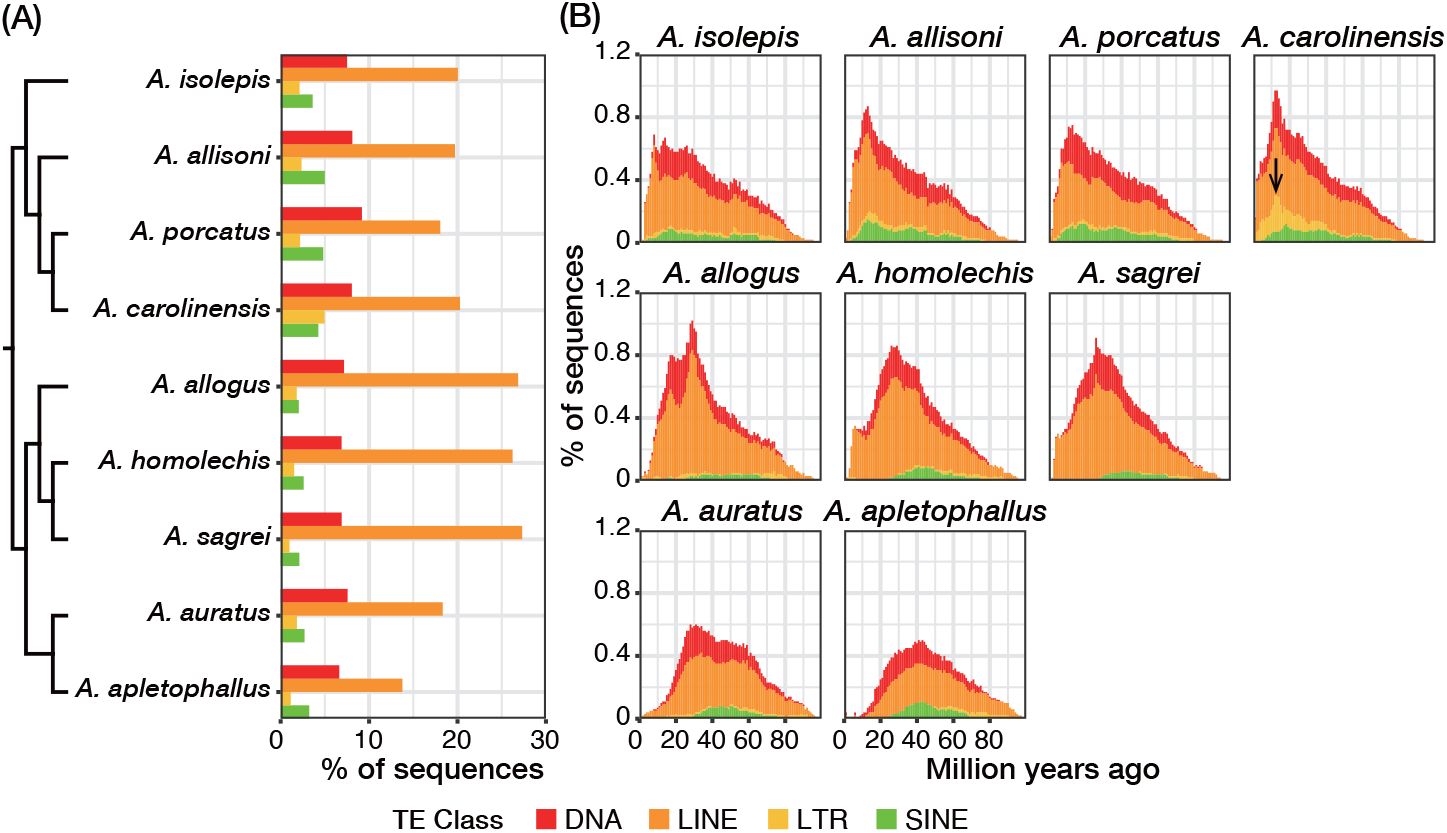
Repeat content and repeat landscape of *Anolis* genomes. (A) Phylogenetic relationships between *Anolis* lizards included in the repeat element analysis and the sequence percentages for each transposable element (TE) class. (B) Repeat landscape for each TE class of *Anolis* lizards included in the repeat element analysis. The arrow points a steep peak for LTR observed only in *A. carolinensis*.

### Identification and analysis of duplicated genes

The *P*_D_ for *A. isolepis, A. allisoni, A. porcatus, A. allogus, A. homolechis*, and *A. sagrei* was 30.8%–39.4% (Fig. 4; Table 3). G-test of independence of species and the number of duplicated and single genes with STP detected associations between species and the number of duplicated and single genes (*P*-values: 8.2 × 10^−45^ for Cuban trunk-crown species and 2.1 × 10^−18^ for Cuban trunk-ground species). Furthermore, the test detected significant bounds on the ratio of duplicated gene numbers to single gene numbers between *A. isolepis* and *A. allisoni*, and between *A. allisoni* and *A. porcatus* within the Cuban trunk-crown species (*P*-values: 0.014 and 1.2 × 10^−25^, respectively), and between *A. allogus* and *A. sagrei*, and between *A. sagrei* and *A. homolechis* within the Cuban trunk-ground species (*P*-values: 5.2 × 10^−7^ and 0.0021, respectively) (Table 3).

**Table 3.**
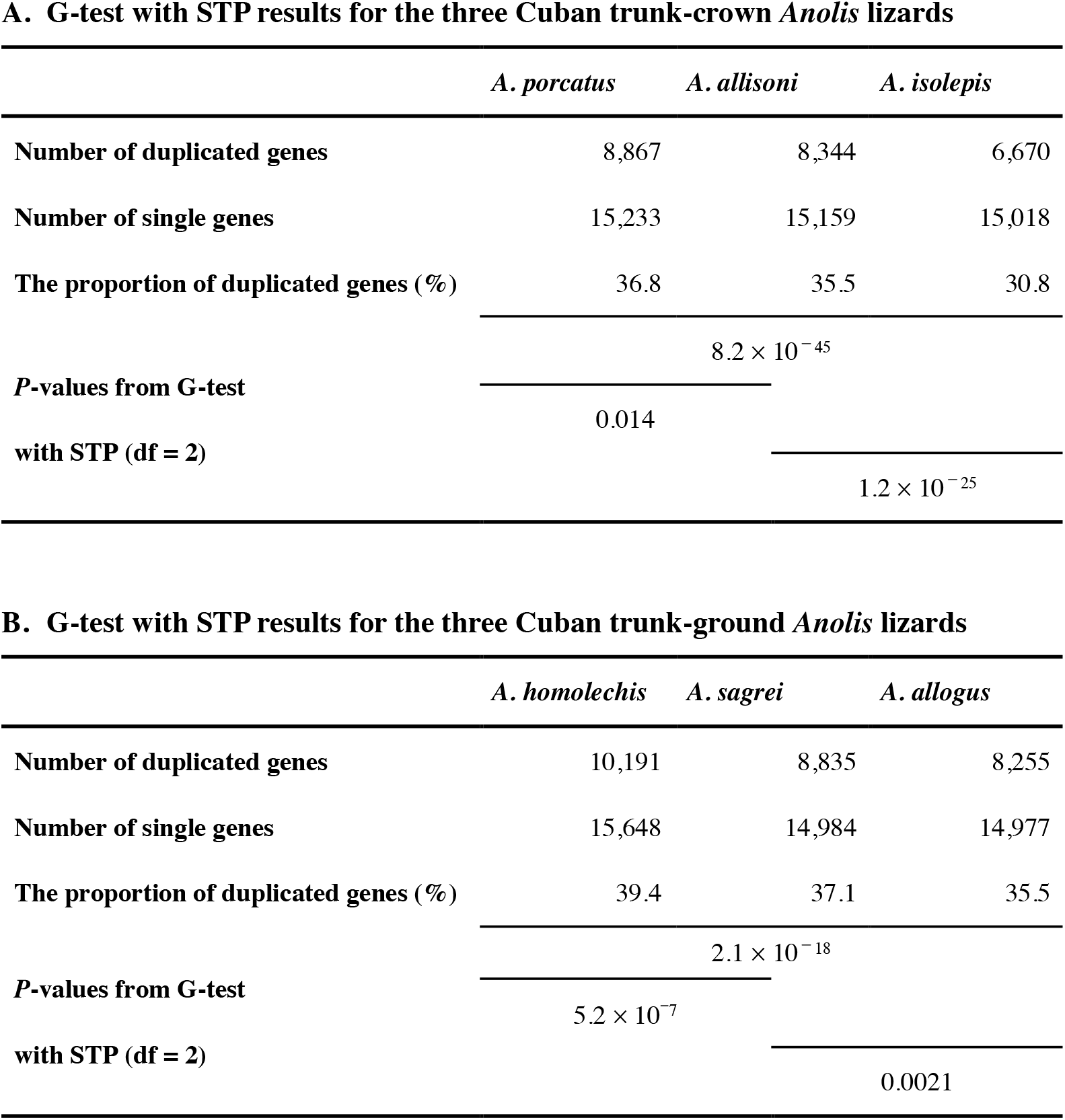
G-test with simultaneous test procedure (STP) results for the homogeneity of duplicated gene to single gene number ratio among (A) three Cuban *Anolis* lizards of the trunk-crown lineage and (B) three Cuban *Anolis* lizards of the trunk-ground lineage. P-values are from the results of G-test with the degree of freedom (df) adjusted to 2 according to simultaneous test procedure (STP, Sokal and Rohlf, 1994) performed among species, which are indicated by black line.

**Fig. 4.**
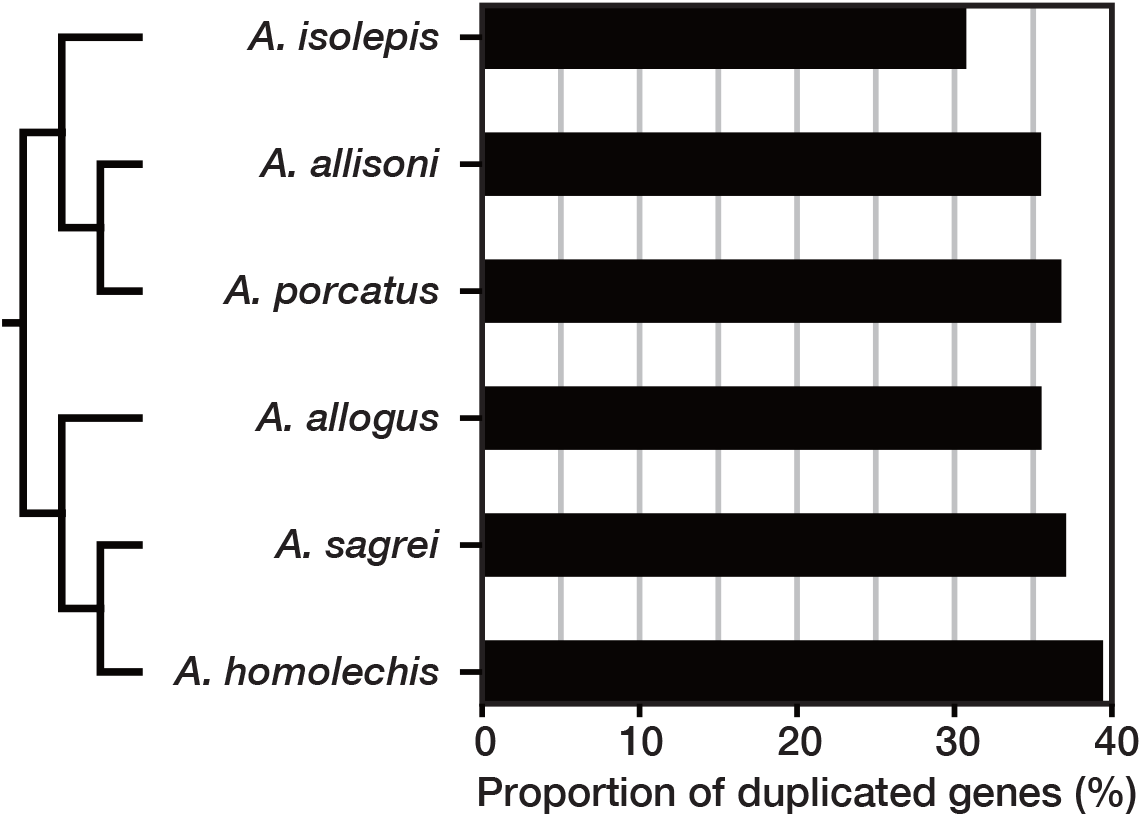
Phylogenetic relationships and proportion of duplicated genes for the six Cuban *Anolis* lizards.

### Estimation of the DNA substitution rate and divergence time

DNA substitution rates (substitutions per a year) at the 4D sites for the branches of the *Anolis* lizards clade in the phylogeny estimated using Bayesian method implemented in MCMCTree ranged from 1.13 × 10^−9^ to 2.20 × 10^−9^ (Supplementary Fig. 3) with an average of 1.8 × 10^−9^. The age of the common ancestor of *Anolis* lizards included in this study and the divergence times of Cuban *Anolis* lizards were estimated to be 51.7 mya in the Eocene and 7.71–38.5 mya from Eocene to Miocene, respectively (Supplementary Fig. 4).

### Estimation of population size history

To estimate the population size history of the six Cuban *Anolis* lizards (*A. isolepis, A. allisoni, A. porcatus, A. allogus, A. homolechis*, and *A. sagrei*), we estimated the past effective population size for each species using PSMC based on the results from mapping back the sequence reads to genome assemblies and calling of the heterozygous sites. Assuming a generation time of one year and mutation rate for *A. isolepis, A. allisoni, A. porcatus, A. allogus, A. homolechis*, and *A. sagrei* of 1.4 × 10^−9^, 1.5 × 10^−9^, 1.8 × 10^−9^, 1.6 × 10^−9^, 2.0 × 10^−9^, and 2.1 × 10^−9^, respectively, the history of effective population size since Pliocene or the latest Miocene, when the divergence of species included in this study was estimated to have generally already ended, was estimated for each species (Fig. 5; Supplementary Fig. 5). An increase in effective population size was estimated for all six species. Furthermore, a significant drop in effective population size was estimated to have occurred in the Middle Pleistocene for *A. porcatus, A. allogus*, and *A. sagrei*. For *A. isolepis*, there was a stagnation in the increase in effective population size during that period. However, for *A. allisoni* and *A. homolechis*, such changes were less pronounced or later than for other species. Change processes in the effective population size of those species did not appear to be linked explicitly to global temperature (Hansen et al. 2013) (Fig. 5).

**Fig. 5.**
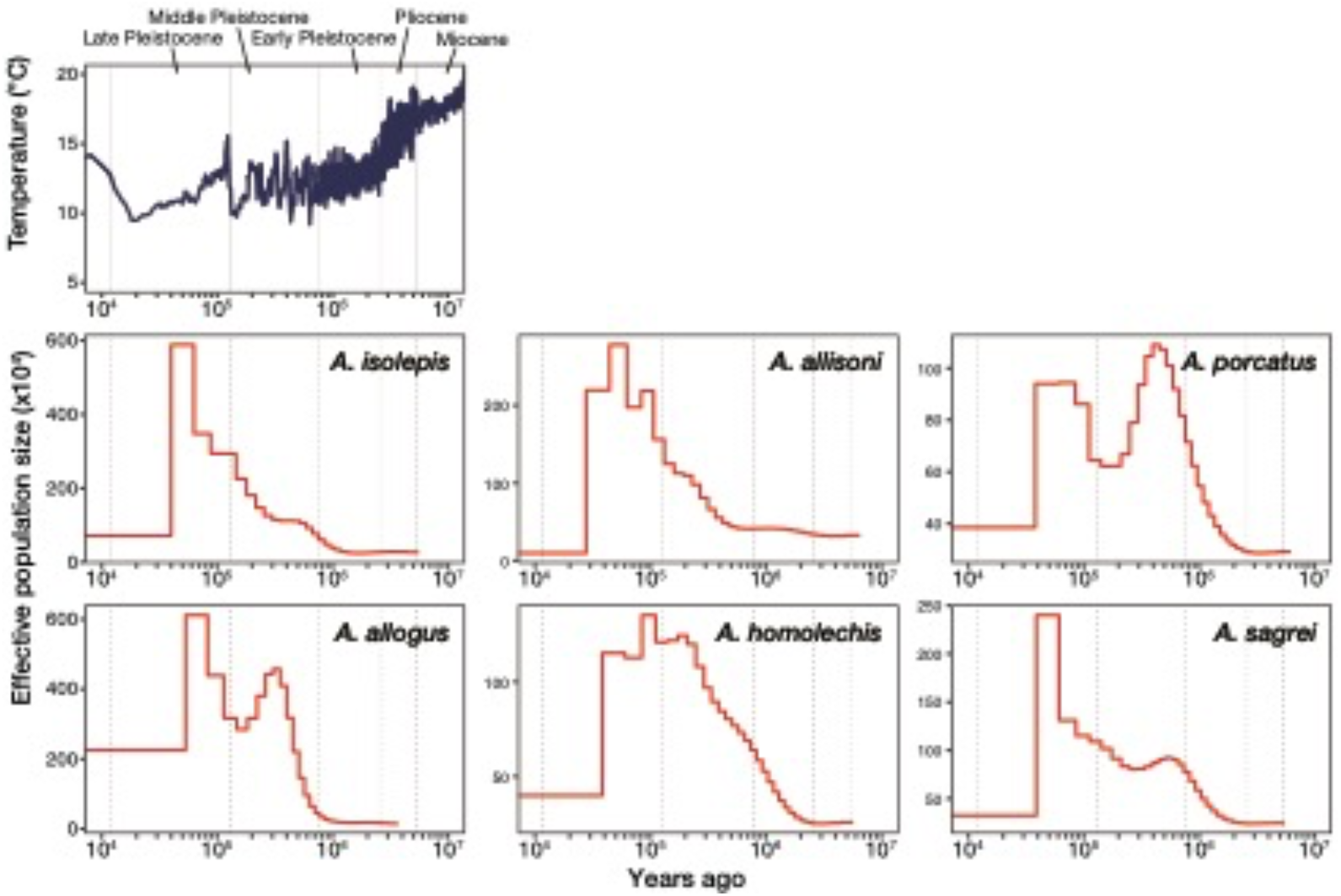
Past effective population sizes of the six Cuban *Anolis* lizards. Top: the earth’s surface temperature (Hansen et al. 2013); six figures below: the estimated past effective population sizes of six Cuban *Anolis* lizards estimated using PSMC. A generation time was set to be one year for all six species. The mutation rate for *A. isolepis, A. allisoni, A. porcatus, A. allogus, A. homolechis*, and *A. sagrei* was set to be 1.4 × 10^−9^, 1.5 × 10^−9^, 1.8 × 10^−9^, 1.6 × 10^−9^, 2.0 × 10^−9^, and 2.1 × 10^−9^, respectively.

## Discussion

We reported the genomes of six Cuban *Anolis* lizards (*A. isolepis, A. allisoni, A. porcatus, A. allogus, A. homolechis*, and *A. sagrei*), three each from trunk-crown (*A. isolepis, A. allisoni*, and *A. porcatus*) and trunk-ground (*A. allogus, A. homolechis*, and *A. sagrei*) lineages of Cuban *Anolis* lizards, which also varied in habitats. These genome assemblies provide genomic resources for the elucidating genetic basis of the diversification and adaptive evolution. In this study, by comparative analysis of these genomes with the genomes of mainland *Anolis* lizards reported in previous studies, we attempted to elucidate the creation and accumulation process of genetic variation during the diversification of *Anolis* lizards.

The coverage of completed BUSCOs was high (77.5%–86.9%) in the six novel genome assemblies reconstructed in this study, comparable to that of the *A. carolinensis* genome (AnoCar2.0) (88.4%), which was assembled to the chromosome level (Fig.1; Supplementary Table 4). Furthermore, the number of genes predicted for each species was 21,688–25,839, similar to that for *A. carolinensis* registered in Ensembl release 104 (21,555). Thus, our obtained novel draft genomes had high gene coverage.

GC content appeared to be slightly higher in the three Cuban trunk-ground *Anolis* lizards than in the three Cuban trunk-crown *Anolis* lizards and their relative, *A. carolinensis* (Fig. 2-1). Furthermore, the shape of GC distribution appeared to be a difference between the three mainland species (*A. frenatus, A. auratus, A. apletophallus*) and six Cuban species and their relative, *A. carolinensis* (Fig. 2). We also observed that the GC distribution of six Cuban species and their relative, *A. carolinensis*, are more symmetrical and steeper. GC content is affected by DNA methylation and recombination rates (Mugal et al. 2015). Since altering the DNA methylation site could results in change of gene expression regulation dynamics and recombination, it is important source for genomic evolution. Hence, differences of GC distribution among lineages or species may reflect different process of genomic adaptive evolution. However, it should be noted that the assembly sizes of novel draft genomes of six Cuban *Anolis* lizards reconstructed in this study were smaller than their estimated genome sizes and still had many gaps (5.6%–9.3% of genomes). Previously, Costantini et al. (2016) also argued that the *A. carolinensis* genome had many gaps due to high GC content, which makes sequencing difficult. Therefore, whether the difference in GC content and distribution among lineages or species is related to the evolution of the location and amount of DNA methylation site and recombination site in genomes is a question for future work. This investigation requires a comparison of the GC landscape at the chromosome level of genome assembly with little gap for many species.

The average of estimated DNA substitution rates (substitutions per a year) for *Anolis* lizard branches was 1.8 × 10^−9^, close to that calculated for mainland *Anolis* lizards by Tollis et al. (2018). Estimated age of the common ancestor of *Anolis* lizards included in this study, 51.7 mya (Supplementary Fig. 4), was roughly consistent with previous studies (Poe et al. 2017; Román-palacios et al. 2018; Tollis et al. 2018). Estimated divergence times of Cuban *Anolis* lizards, 7.71–38.5 mya, (Supplementary Fig. 4) were also roughly consistent with the estimation of previous studies (Poe et al. 2017; Román-Palacios et al. 2018). However, divergence time of Cuban *Anolis* lizards is slightly more recent than those in previous estimates. Although considerably more genes and sites were used in the analysis than in previous studies, it should be noted that the results of this study are not entirely correct either, since functional constraints were observed in the 4D site (Künstner et al. 2011), which we used for the analysis.

Some differences in the composition of repeat elements were also observed among species. By comparing the composition of repeat elements and repeat landscape for six Cuban *Anolis* lizards plus three mainland *Anolis* lizards (*A. carolinensis, A. auratus*, and *A. apletophallus*), with the phylogenetic relationships and divergence time of their species, an overview of repeat element accumulation process during the diversification of *Anolis* lizards could be provided. The genome of *A. carolinensis* contains the largest amount of LTR repeats (Fig. 3; Supplementary Tables 5–13) and only its repeat landscape showed a steep LTR repeat accumulation wave that peaked around 10 mya, mainly composed of the Gypsy family (Supplementary Fig. 1). Although it was already noted that the genome of *A. carolinensis* genome had more LTRs than that of *A. sagrei* (Geneva et al. 2021), we further revealed that it had considerably more LTRs than six Cuban *Anolis* lizards, including three closely related Cuban trunk-crown species (*A. isolepis, A. allisoni*, and *A. porcatus*). We also observed that the steep wave of LTRs in the repeat landscape for *A. carolinensis* was lacking in those for these six Cuban species. Since the divergence time between *A. carolinensis* and *A. allisoni*, and *A. carolinensis* and *A. isolepis* was estimated to be 7.71 and 14.03 mya, respectively, the LTRs amplified in the common ancestor of *A. carolinensis* and *A. allisoni* might have accumulated only in the ancestors of *A. carolinensis*. Additionally, looking at the overall shape of the repeat landscapes, those of the trunk-crown Cuban *Anolis* lizards and their relative, *A. carolinensis* had more recent peaks than those of other species, and the age of the peak for closely related *A. auratus* and *A. apletophallus* were similar to those of trunk-ground Cuban *Anolis* lizards, although the shape were steeper in trunk-ground Cuban *Anolis* lizards. These results indicate that each lineage had undergone different accumulation processes of repeat elements. Results also showed that the genomes of three Cuban *Anolis* lizards contained more LINEs than other *Anolis* lizards (Fig. 3; Supplementary Table 5 to 13). However, comparing of their breakdown (Supplementary Table 5 to 13) and repeat landscapes (Supplementary Fig. 2), differences in accumulation processes of several LINE families were suggested among these species. Therefore, it is possible that their common ancestor experienced changes in something related to common transposition mechanisms among the LINE families.

Significant differences in the ratio of duplicated genes to single copy genes were detected among Cuban *Anolis* lizards in both trunk-crown and trunk-ground lineages. Results indicated that forest species; *A. isolepis* and *A. allogus* had the lowest *P*_D_ than open areas and forest margin species in each lineage. Notably, *A. isolepis*, which lives only in mountainous cloud forests or rain forests at more than 800 m above sea level (Schettino 1999), had the lowest *P*_D_. We also detected significant differences in the ratio of duplicated genes to single copy genes between *A. porcatus* and *A. allisoni* inhabiting open areas in the trunk-crown lineage, and between *A. homolechis* and *A. sagrei* inhabiting forest margin and open areas, respectively, in trunk-ground lineage. *A. homolechis*, which showed the highest *P*_D_, have a wider range of body temperatures in natural area of both males and females than other species (Schettino et al. 2010; Ruibal 1961). Gene duplication can be a potent source of variation that is required for adaptive evolution. Makino and Kawata (2012) and Tamate et al. (2014) showed that the range of habitat climate is positively correlated with *P*_D_. Forests can buffer the climate (e.g., De Frenne et al. 2019; Lin et al. 2020). Therefore, it is conceivable that the range of climatic conditions was narrower in the forest than in open areas and forest edges, and that it may be related to *P*_D_ of Cuban *Anolis* lizards. Makino and Kawata (2019) also showed that invasive species have high *P*_D_. *A. porcatus* is a close relative of *A. carolinensis*, which is native to the United States and has invaded many parts of the world. The higher *P*_D_ in *A. porcatus* than in *A. allisoni* might be related to the potential invasiveness of *A. porcatus*. Given that the evolution of new copies created by duplication may facilitate advancing into novel environments, it is also possible that the direction of evolution, i.e., whether the habitat is ancestral or not, is related to *P*_D_. Thus, to verify the relationship between *P*_D_ and natural habitats more accurately, it will be necessary to examine the habitat transition process and analyze *P*_D_ with more species.

Estimating population size history can provide important information for examining the evolutionary process of organisms. We estimated past effective population sizes of six Cuban *Anolis* lizards since the Pliocene or the latest Miocene (Fig. 5). If the mutation rate and generation time set in this study are correct, the following can be inferred from the results. Although an increase in effective population size was estimated for all six species, for *A. isolepis, A. porcatus, A. allogus*, and *A. sagrei*, a fall or cessation of the increase in the Middle Pleistocene was observed. However, for *A. allisoni* and *A. homolechis*, such changes were less pronounced or appeared later than in the other species. Paleogeographic studies proposed that, although high altitudes in parts of central and eastern Cuba were not inundated, many parts of Cuba were inundated in the sea near the boundary between the Middle and Late Pleistocene, compared to the Pliocene and early Pleistocene periods and present-day (Iturralde-Vinent 2003, 2006). Individuals of *A. isolepis, A. allogus*, and *A. sagrei*, whose genomes were analyzed in this study, were collected from western Cuba. Therefore, we consider that the fluctuation in habitat range due to inundation into the sea might have contributed to the population size fluctuation of these species in western Cuba. The smaller fluctuation for *A. isolepis* compared to *A. allogus* and *A. sagrei* may be because the distribution of *A. isolepis* was limited to high elevation mountainous areas (Schettino 1999). An individual of *A. homolechis* was collected from eastern Cuba. It is also assumed that the effect of fluctuations in sea level on population size was smaller or occurred later in eastern Cuba than in western Cuba. *A. porcatus* and *A. allisoni* samples were collected in the central part of the country, but for *A. porcatus*, severe fluctuation in effective population size was estimated. *A. porcatus* and *A. allisoni* are closely related, with larger size of *A. allisoni* currently dominating in central Cuba (Schettino 1999; Glor et al. 2004). Since the beginning of the decline in the effective population size of *A. porcatus* and an increase in the effective population size of *A. allisoni* roughly overlapped in the Middle Pleistocene, the pattern of fluctuations in the effective population sizes of populations in central Cuba of these two species may reflect an intense interspecific competition. Moreover, an rise in effective population size of *A. isolepis* and *A. allogus*, which are currently inhabiting forests, were estimated around 1 million years ago, while that of *A. porcatus, A. sagrei*, and *A. homolechis*, which are currently inhabiting open areas or forest margins other than *A. allisoni*, were estimated a little earlier, around 2 million years ago. How long the current habitat of each species have remained the same is unknown, but if they have remained the same since 2 million years ago, such differences may be due to differences in habitats. Thus, if the time scale of the estimation is correct, it is conceivable that the history of population sizes of these six Cuban *Anolis* lizards is related to Cuba’s geohistory, interspecific competition, and habitats.

In this study, we reconstructed novel genome assemblies of six Cuban *Anolis* lizards with relatively long and high gene completeness. Then, by examining and comparing each feature of the genome, including those of mainland species reported in previous studies, we estimated the genetic variation that occurred during the diversification of *Anolis* lizards. Interspecific comparisons for the TE and gene composition detected the different TE accumulation process and gene copy number change for each species or lineage. Furthermore, estimates of the past effective population size suggest the possibility that population size of these species may have fluctuated due to geohistory or interspecific competition. These results are expected to provide important information as a stepping stone for elucidating the diversification and adaptive radiation mechanism of *Anolis* lizards. Further improvements in the quality of the novel genome assemblies reconstructed in this study are expected to advance population and epigenome analyses at the genome level.

## Acknowledgments

We are grateful to Hisayo Asao and Dr. Asaka Akita at the NIBB Core Research Facilities, National Institute for Basic Biology for DNA extraction and sequence library preparation. We also thank Anthony Geneva and Jonathan Losos for commenting on our manuscript. Computations were partially performed on the NIG supercomputer at ROIS National Institute of Genetics. This work was supported by a Grant-in-Aid for Scientific Research(16H05767 and 19KK0184) from the Japan Society for the Promotion of Science, HFSP research grant (RGP0030/2020) and NIBB Collaborative Research Program (18-403) to MK.

## Data availability

The paired-end reads, which were used for *de novo* genome assembly, were deposited in the DDBJ Short Read Archive database. The set of the soft-masked genome assemblies, gene models, and predicted coding nucleotide and peptide sequences are available at figshare.

## Contributions

SK and MK designed the study. SK, LMD, AC, and MK collected samples. KY and SS directed library preparation with the Chromium system and sequencing. SK reconstructed the draft genome assemblies and performed all sequence analyses. SK and MK wrote the manuscript. All authors read and approved the final manuscript.

