## Supplemental Tables and Figures for "Draft genome of six Cuban *Anolis* lizards and insights into genetic changes during the diversification"

**Supplementary Table 1. Run IDs and accession numbers of RNA-seq data used for training of gene prediction on genome assemblies in the Sequence Read Archive of the DNA Data Bank of Japan (DDBJ).**

| Species | Run IDs in the SRA of the DDBJ | Accession numbers in the SRA of the DDBJ |
|---|---|---|
| *Anolis isolepis* | DRR232283 | DRA010304 |
| *Anolis allisoni* | DRR232284 | DRA010304 |
| *Anolis porcatus* | DRR232285 | DRA010304 |
| *Anolis allogus* | DRR055059, DRR055051, DRR055067 | DRA004457 |
| *Anolis homolechis* | DRR055075, DRR055083, DRR055091 | DRA004457 |
| *Anolis sagrei* | DRR055099, DRR055107, DRR055115 | DRA004457 |

**Supplementary Table 2. Source database of sequence or genome annotation data used in phylogenetic analysis.**

| Species | Genome assembly | The source database of sequence FASTA files and/or GFF file |
|---|---|---|
| *Anas platyrhynchos* | CAU_duck1.0 | Ensemble (release 104) |
| *Anolis apletophallus* | Aapl1.0 | Harvard Dataverse |
| *Anolis auratus* | Aaur1.0 | Harvard Dataverse |
| *Anolis carolinensis* | AnoCar2.0 | Ensemble (release 104) |
| *Anolis frenatus* | Afre1.0 | Harvard Dataverse |
| *Aquila chrysaetos* | bAquChr1.2 | Ensemble (release 104) |
| *Bos taurus* | UOA_Angus_1 | Ensemble (release 104) |
| *Canis lupus* | CanFam3.1 | Ensemble (release 104) |
| *Chelonoidis abingdonii* | ASM359739v1 | Ensemble (release 104) |
| *Chrysemys picta* | Chrysemys_picta_bellii-3.0.3 | Ensemble (release 104) |
| *Crocodylus porosus* | CroPor_comp1 | Ensemble (release 104) |
| *Felis catus* | Felis_catus_9.0 | Ensemble (release 104) |
| *Ficedula albicollis* | FicAlb1.5 | Ensemble (release 104) |
| *Gallus gallus* | GRCg6a | Ensemble (release 104) |
| *Gekko japonicus* | Gekko_japonicus_V1.1 | NCBI |
| *Geospiza fortis* | GeoFor_1.0 | Ensemble (release 104) |
| *Gopherus evgoodei* | rGopEvg1_v1.p | Ensemble (release 104) |
| *Homo sapiens* | GRCh38.p13 | Ensemble (release 104) |
| *Laticauda laticaudata* | latLat_1.0 | Ensemble (release 104) |
| *Latimeria chalumnae* | LatCha1 | Ensemble (release 104) |
| *Meleagris gallopavo* | Turkey_5.1 | Ensemble (release 104) |
| *Monodelphis domestica* | ASM229v1 | Ensemble (release 104) |
| *Mus musculus* | GRCm39 | Ensemble (release 104) |
| *Myotis lucifugus* | Myoluc2.0 | Ensemble (release 104) |
| *Naja naja* | Nana_v5 | Ensemble (release 104) |
| *Notechis scutatus* | TS10Xv2-PRI | Ensemble (release 104) |
| *Ornithorhynchus anatinus* | mOrnAna1.p.v1 | Ensemble (release 104) |
| *Pelodiscus sinensis* | PelSin_1.0 | Ensemble (release 104) |

| | | |
|---|---|---|
| *Pogona vitticeps* | pvi1.1 | Ensemble (release 104) |
| *Pseudonaja textilis* | EBS10Xv2-PRI | Ensemble (release 104) |
| *Struthio camelus* | ASM69896v1 | Ensemble (release 104) |
| *Sus scrofa* | Sscrofa11.1 | Ensemble (release 104) |
| *Taeniopygia guttata* | bTaeGut1_v1.p | Ensemble (release 104) |
| *Terrapene carolina* | T_m_triunguis-2.0 | Ensemble (release 104) |
| *Tursiops truncatus* | turTru1 | Ensemble (release 104) |
| *Ursus maritimus* | UrsMar_1.0 | Ensemble (release 104) |

**Supplementary Table 3. Genome sequencing and *de novo* genome assembly results before removing haplotigs and overlaps.**

| Species | Total read data before adjusting the number of reads (Gb) | Coverage before adjusting the number of reads | Coverage after adjusting the number of reads | Contig N50 (Kb) | Scaffold N50 (Mb) | Total length (Gb) | Number ≧ 10Kb |
|---|---|---|---|---|---|---|---|
| *Anolis isolepis* | 128 | 61.70× | 56.64× | 62.59 | 28.69 | 1.53 | 3.36 K |
| *Anolis allisoni* | 129 | 57.34× | 56.27× | 43.31 | 5.69 | 1.57 | 7.31 K |
| *Anolis porcatus* | 134 | 63.30× | 56.80× | 49.41 | 22.02 | 1.55 | 7.90 K |
| *Anolis allogus* | 125 | 54.16× | | 46.44 | 42.71 | 1.88 | 6.61 K |
| *Anolis homolechis* | 125 | 47.26× | | 44.47 | 41.02 | 1.65 | 11.46 K |
| *Anolis sagrei* | 123 | 60.18× | 56.48× | 49.79 | 34.06 | 1.72 | 11.18 K |

**Supplementary Table 4. BUSCO metrics of genome assemblies.** BUSCO: Benchmarking Universal Single-Copy Orthologs

| | **Before purge of haplotigs and overlaps** | | | | **After purge of haplotigs and overlaps** | | | |
|---|---|---|---|---|---|---|---|---|
| **Species** | **Complete** | **(Single** | **Duplicated)** | **Fragmented** | **Complete** | **(Single** | **Duplicated)** | **Fragmented** |
| *Anolis isolepis* | 88.0% | 86.2% | 1.8% | 7.7% | 86.9% | 86.1% | 0.8% | 7.0% |
| *Anolis allisoni* | 85.9% | 84.2% | 1.7% | 9.6% | 83.8% | 83.1% | 0.7% | 9.4% |
| *Anolis porcatus* | 85.7% | 83.4% | 2.3% | 10.0% | 83.7% | 83.1% | 0.6% | 9.4% |
| *Anolis allogus* | 86.7% | 82.4% | 4.3% | 8.5% | 86.0% | 83.8% | 2.2% | 8.4% |
| *Anolis homolechis* | 79.4% | 77.8% | 1.6% | 12.8% | 77.5% | 76.6% | 0.9% | 12.3% |
| *Anolis sagrei* | 86.0% | 83.6% | 2.4% | 9.4% | 85.0% | 84.0% | 1.0% | 8.9% |

**Supplementary Table 5. Content of repeat elements in the *Anolis isolepis* genome.**

| | Number of elements | Length occupied (bp) | Percentage of sequence |
|---|---|---|---|
| **Retroelements** | 1878168 | 430006788 | 25.81 |
| **SINEs:** | 368401 | 60380616 | 3.62 |
| **Penelope** | 324880 | 45110106 | 2.71 |
| **LINEs:** | 1430507 | 333966228 | 20.04 |
| **CRE/SLACS** | 0 | 0 | 0 |
| **L2/CR1/Rex** | 770129 | 166127355 | 9.97 |
| **R1/LOA/Jockey** | 0 | 0 | 0 |
| **R2/R4/NeSL** | 38810 | 13264806 | 0.8 |
| **RTE/Bov-B** | 191555 | 56599760 | 3.4 |
| **L1/CIN4** | 61011 | 35495653 | 2.13 |
| **LTR elements:** | 79260 | 35659944 | 2.14 |
| **BEL/Pao** | 1374 | 861407 | 0.05 |
| **Ty1/Copia** | 25897 | 5566259 | 0.33 |
| **Gypsy/DIRS1** | 34004 | 22064860 | 1.32 |
| **Retroviral** | 11690 | 4262542 | 0.26 |
| **DNA transposons** | 815039 | 125105108 | 7.51 |
| **hobo-Activator** | 304052 | 42543725 | 2.55 |
| **Tc1-IS630-Pogo** | 341448 | 61163483 | 3.67 |
| **En-Spm** | 0 | 0 | 0 |
| **MuDR-IS905** | 0 | 0 | 0 |
| **PiggyBac** | 0 | 0 | 0 |
| **Tourist/Harbinger** | 124257 | 15460963 | 0.93 |
| **Other** | 0 | 0 | 0 |
| **Rolling-circles** | 103984 | 27359258 | 1.64 |
| **Unclassified:** | 857 | 132814 | 0.01 |
| **Total interspersed repeats:** | | 555244710 | 33.32 |
| **Small RNA:** | 3962 | 421256 | 0.03 |
| **Satellites:** | 4873 | 1133691 | 0.07 |
| **Simple repeats:** | 464709 | 19405729 | 1.16 |
| **Low complexity:** | 42891 | 2546122 | 0.15 |

**Supplementary Table 6. Content of repeat elements in the *Anolis allisoni* genome.**

| | Number of elements | Length occupied (bp) | Percentage of sequence |
|---|---|---|---|
| **Retroelements** | 1998283 | 472471996 | 27.06 |
| **SINEs:** | 493079 | 87160708 | 4.99 |
| **Penelope** | 200757 | 36629163 | 2.1 |
| **LINEs:** | 1416121 | 344051638 | 19.71 |
| **CRE/SLACS** | 0 | 0 | 0 |
| **L2/CR1/Rex** | 808487 | 177785673 | 10.18 |
| **R1/LOA/Jockey** | 0 | 0 | 0 |
| **R2/R4/NeSL** | 96899 | 33309694 | 1.91 |
| **RTE/Bov-B** | 200522 | 50301566 | 2.88 |
| **L1/CIN4** | 71273 | 33018779 | 1.89 |
| **LTR elements:** | 89083 | 41259650 | 2.36 |
| **BEL/Pao** | 2511 | 1608912 | 0.09 |
| **Ty1/Copia** | 13499 | 5445708 | 0.31 |
| **Gypsy/DIRS1** | 61792 | 28108497 | 1.61 |
| **Retroviral** | 7992 | 4342287 | 0.25 |
| **DNA transposons** | 935292 | 141605863 | 8.11 |
| **hobo-Activator** | 392108 | 59692810 | 3.42 |
| **Tc1-IS630-Pogo** | 349042 | 57717916 | 3.31 |
| **En-Spm** | 0 | 0 | 0 |
| **MuDR-IS905** | 0 | 0 | 0 |
| **PiggyBac** | 362 | 93855 | 0.01 |
| **Tourist/Harbinger** | 123305 | 14740173 | 0.84 |
| **Other** | 0 | 0 | 0 |
| **Rolling-circles** | 90446 | 15765657 | 0.9 |
| **Unclassified:** | 847 | 130180 | 0.01 |
| **Total interspersed repeats:** | na | 614208039 | 35.18 |
| **Small RNA:** | 29050 | 3533512 | 0.2 |
| **Satellites:** | 5810 | 1242447 | 0.07 |
| **Simple repeats:** | 523272 | 21575630 | 1.24 |
| **Low complexity:** | 43639 | 2563403 | 0.15 |

**Supplementary Table 7. Content of repeat elements in the *Anolis porcatus* genome.**

| | Number of elements | Length occupied (bp) | Percentage of sequence |
|---|---|---|---|
| **Retroelements** | 1930718 | 436494158 | 25.05 |
| **SINEs:** | 461468 | 83754369 | 4.81 |
| **Penelope** | 246114 | 39263710 | 2.25 |
| **LINEs:** | 1365220 | 314491428 | 18.05 |
| **CRE/SLACS** | 0 | 0 | 0 |
| **L2/CR1/Rex** | 647860 | 142289738 | 8.17 |
| **R1/LOA/Jockey** | 0 | 0 | 0 |
| **R2/R4/NeSL** | 82081 | 26647143 | 1.53 |
| **RTE/Bov-B** | 184598 | 47877138 | 2.75 |
| **L1/CIN4** | 53402 | 27608041 | 1.58 |
| **LTR elements:** | 104030 | 38248361 | 2.2 |
| **BEL/Pao** | 2372 | 1477202 | 0.08 |
| **Ty1/Copia** | 13214 | 5295606 | 0.3 |
| **Gypsy/DIRS1** | 75415 | 26203429 | 1.5 |
| **Retroviral** | 5210 | 3173081 | 0.18 |
| **DNA transposons** | 1050505 | 160076517 | 9.19 |
| **hobo-Activator** | 572817 | 80050700 | 4.59 |
| **Tc1-IS630-Pogo** | 344059 | 63690746 | 3.66 |
| **En-Spm** | 0 | 0 | 0 |
| **MuDR-IS905** | 0 | 0 | 0 |
| **PiggyBac** | 331 | 91451 | 0.01 |
| **Tourist/Harbinger** | 95172 | 11036882 | 0.63 |
| **Other** | 0 | 0 | 0 |
| **Rolling-circles** | 91492 | 19842974 | 1.14 |
| **Unclassified:** | 854 | 130249 | 0.01 |
| **Total interspersed repeats:** | na | 596700924 | 34.24 |
| **Small RNA:** | 3977 | 314281 | 0.02 |
| **Satellites:** | 8895 | 1547840 | 0.09 |
| **Simple repeats:** | 541400 | 23108516 | 1.33 |
| **Low complexity:** | 44267 | 2558821 | 0.15 |

**Supplementary Table 8. Content of repeat elements in the *Anolis allogus* genome.**

| | Number of elements | Length occupied (bp) | Percentage of sequence |
|---|---|---|---|
| **Retroelements** | 2662359 | 642619412 | 30.72 |
| **SINEs:** | 318314 | 42786880 | 2.05 |
| **Penelope** | 176743 | 37730635 | 1.8 |
| **LINEs:** | 2241071 | 561848287 | 26.86 |
| **CRE/SLACS** | 0 | 0 | 0 |
| **L2/CR1/Rex** | 1294566 | 298470694 | 14.27 |
| **R1/LOA/Jockey** | 1668 | 459265 | 0.02 |
| **R2/R4/NeSL** | 35650 | 12561606 | 0.6 |
| **RTE/Bov-B** | 303421 | 76898337 | 3.68 |
| **L1/CIN4** | 69816 | 39473943 | 1.89 |
| **LTR elements:** | 102974 | 37984245 | 1.82 |
| **BEL/Pao** | 28800 | 8259095 | 0.39 |
| **Ty1/Copia** | 18194 | 5559362 | 0.27 |
| **Gypsy/DIRS1** | 30908 | 17969621 | 0.86 |
| **Retroviral** | 16895 | 3008300 | 0.14 |
| **DNA transposons** | 903077 | 149896799 | 7.17 |
| **hobo-Activator** | 439389 | 77691128 | 3.71 |
| **Tc1-IS630-Pogo** | 283381 | 50504196 | 2.41 |
| **En-Spm** | 0 | 0 | 0 |
| **MuDR-IS905** | 0 | 0 | 0 |
| **PiggyBac** | 282 | 62278 | 0 |
| **Tourist/Harbinger** | 129608 | 15453741 | 0.74 |
| **Other** | 0 | 0 | 0 |
| **Rolling-circles** | 133030 | 35287662 | 1.69 |
| **Unclassified:** | 897 | 139208 | 0.01 |
| **Total interspersed repeats:** | na | 792655419 | 37.89 |
| **Small RNA:** | 9454 | 679118 | 0.03 |
| **Satellites:** | 1593 | 145840 | 0.01 |
| **Simple repeats:** | 630240 | 32035685 | 1.53 |
| **Low complexity:** | 63954 | 6340749 | 0.3 |

**Supplementary Table 9. Content of repeat elements in the *Anolis homolechis* genome.**

| | Number of elements | Length occupied (bp) | Percentage of sequence |
|---|---|---|---|
| **Retroelements** | 2343675 | 597838716 | 30.37 |
| **SINEs:** | 305541 | 51252321 | 2.6 |
| **Penelope** | 215376 | 44338146 | 2.25 |
| **LINEs:** | 1971309 | 516421231 | 26.23 |
| **CRE/SLACS** | 0 | 0 | 0 |
| **L2/CR1/Rex** | 1158109 | 287403950 | 14.6 |
| **R1/LOA/Jockey** | 0 | 0 | 0 |
| **R2/R4/NeSL** | 53553 | 13242368 | 0.67 |
| **RTE/Bov-B** | 320262 | 84847075 | 4.31 |
| **L1/CIN4** | 52083 | 28870316 | 1.47 |
| **LTR elements:** | 66825 | 30165164 | 1.53 |
| **BEL/Pao** | 12206 | 7709023 | 0.39 |
| **Ty1/Copia** | 5374 | 2218471 | 0.11 |
| **Gypsy/DIRS1** | 33551 | 14359346 | 0.73 |
| **Retroviral** | 2864 | 1255456 | 0.06 |
| **DNA transposons** | 935915 | 135737454 | 6.89 |
| **hobo-Activator** | 513728 | 70880574 | 3.6 |
| **Tc1-IS630-Pogo** | 234336 | 42194176 | 2.14 |
| **En-Spm** | 0 | 0 | 0 |
| **MuDR-IS905** | 0 | 0 | 0 |
| **PiggyBac** | 203 | 47232 | 0 |
| **Tourist/Harbinger** | 138615 | 16355869 | 0.83 |
| **Other** | 0 | 0 | 0 |
| **Rolling-circles** | 147579 | 25177678 | 1.28 |
| **Unclassified:** | 871 | 135886 | 0.01 |
| **Total interspersed repeats:** | na | 733712056 | 37.27 |
| **Small RNA:** | 51342 | 5240052 | 0.27 |
| **Satellites:** | 2555 | 1038946 | 0.05 |
| **Simple repeats:** | 603318 | 27492589 | 1.4 |
| **Low complexity:** | 54762 | 4429680 | 0.23 |

**Supplementary Table 10. Content of repeat elements in the *Anolis sagrei* genome.**

| | Number of elements | Length occupied (bp) | Percentage of sequence |
|---|---|---|---|
| **Retroelements** | 2522210 | 579096787 | 30.43 |
| **SINEs:** | 282067 | 40211670 | 2.11 |
| **Penelope** | 210517 | 40834967 | 2.15 |
| **LINEs:** | 2194773 | 519825794 | 27.31 |
| **CRE/SLACS** | 0 | 0 | 0 |
| **L2/CR1/Rex** | 1296452 | 301555406 | 15.85 |
| **R1/LOA/Jockey** | 0 | 0 | 0 |
| **R2/R4/NeSL** | 46306 | 12524841 | 0.66 |
| **RTE/Bov-B** | 330586 | 83497794 | 4.39 |
| **L1/CIN4** | 49161 | 22657559 | 1.19 |
| **LTR elements:** | 45370 | 19059323 | 1 |
| **BEL/Pao** | 5336 | 2436534 | 0.13 |
| **Ty1/Copia** | 7430 | 3242174 | 0.17 |
| **Gypsy/DIRS1** | 17925 | 9717670 | 0.51 |
| **Retroviral** | 8711 | 1035769 | 0.05 |
| **DNA transposons** | 927191 | 131317724 | 6.9 |
| **hobo-Activator** | 479381 | 66069988 | 3.47 |
| **Tc1-IS630-Pogo** | 272130 | 44239274 | 2.32 |
| **En-Spm** | 0 | 0 | 0 |
| **MuDR-IS905** | 0 | 0 | 0 |
| **PiggyBac** | 0 | 0 | 0 |
| **Tourist/Harbinger** | 134818 | 15560202 | 0.82 |
| **Other** | 0 | 0 | 0 |
| **Rolling-circles** | 160026 | 33123415 | 1.74 |
| **Unclassified:** | 882 | 137231 | 0.01 |
| **Total interspersed repeats:** | na | 710551742 | 37.34 |
| **Small RNA:** | 34462 | 2951726 | 0.16 |
| **Satellites:** | 2953 | 838302 | 0.04 |
| **Simple repeats:** | 580397 | 28765686 | 1.51 |
| **Low complexity:** | 54937 | 4853191 | 0.26 |

**Supplementary Table 11. Content of repeat elements in the *Anolis carolinensis* genome.**

| | Number of elements | Length occupied (bp) | Percentage of sequence |
|---|---|---|---|
| **Retroelements** | 2242332 | 530789914 | 29.5 |
| **SINEs:** | 443355 | 76584647 | 4.26 |
| **Penelope** | 393954 | 57074069 | 3.17 |
| **LINEs:** | 1684725 | 364913695 | 20.28 |
| **CRE/SLACS** | 0 | 0 | 0 |
| **L2/CR1/Rex** | 704649 | 164713219 | 9.16 |
| **R1/LOA/Jockey** | 0 | 0 | 0 |
| **R2/R4/NeSL** | 244595 | 36955059 | 2.05 |
| **RTE/Bov-B** | 223337 | 56970677 | 3.17 |
| **L1/CIN4** | 55123 | 29525681 | 1.64 |
| **LTR elements:** | 114252 | 89291572 | 4.96 |
| **BEL/Pao** | 5173 | 7158627 | 0.4 |
| **Ty1/Copia** | 12320 | 5283274 | 0.29 |
| **Gypsy/DIRS1** | 73032 | 62935021 | 3.5 |
| **Retroviral** | 18936 | 10153865 | 0.56 |
| **DNA transposons** | 861886 | 144912403 | 8.05 |
| **hobo-Activator** | 369164 | 60428226 | 3.36 |
| **Tc1-IS630-Pogo** | 369965 | 68274090 | 3.79 |
| **En-Spm** | 0 | 0 | 0 |
| **MuDR-IS905** | 0 | 0 | 0 |
| **PiggyBac** | 969 | 168940 | 0.01 |
| **Tourist/Harbinger** | 81484 | 9662784 | 0.54 |
| **Other** | 0 | 0 | 0 |
| **Rolling-circles** | 93920 | 24254015 | 1.35 |
| **Unclassified:** | 1506 | 266832 | 0.01 |
| **Total interspersed repeats:** | na | 675969149 | 37.57 |
| **Small RNA:** | 29837 | 2578761 | 0.14 |
| **Satellites:** | 6099 | 1902809 | 0.11 |
| **Simple repeats:** | 492959 | 21208777 | 1.18 |
| **Low complexity:** | 42031 | 2338505 | 0.13 |

**Supplementary Table 12. Content of repeat elements in *Anolis apletophallus* genome.**

| | Number of elements | Length occupied (bp) | Percentage of sequence |
|---|---|---|---|
| **Retroelements** | 1993760 | 395743252 | 18.14 |
| **SINEs:** | 467747 | 70466223 | 3.23 |
| **Penelope** | 182225 | 32773695 | 1.5 |
| **LINEs:** | 1455796 | 300410743 | 13.77 |
| **CRE/SLACS** | 0 | 0 | 0 |
| **L2/CR1/Rex** | 841398 | 162319926 | 7.44 |
| **R1/LOA/Jockey** | 322 | 70651 | 0 |
| **R2/R4/NeSL** | 28895 | 7044990 | 0.32 |
| **RTE/Bov-B** | 261656 | 43587144 | 2 |
| **L1/CIN4** | 92457 | 40869628 | 1.87 |
| **LTR elements:** | 70217 | 24866286 | 1.14 |
| **BEL/Pao** | 3199 | 794646 | 0.04 |
| **Ty1/Copia** | 3479 | 1215599 | 0.06 |
| **Gypsy/DIRS1** | 46237 | 17351451 | 0.8 |
| **Retroviral** | 3198 | 1551805 | 0.07 |
| **DNA transposons** | 1197143 | 144667508 | 6.63 |
| **hobo-Activator** | 674951 | 78820290 | 3.61 |
| **Tc1-IS630-Pogo** | 259669 | 35250873 | 1.62 |
| **En-Spm** | 0 | 0 | 0 |
| **MuDR-IS905** | 0 | 0 | 0 |
| **PiggyBac** | 0 | 0 | 0 |
| **Tourist/Harbinger** | 198545 | 23253126 | 1.07 |
| **Other** | 0 | 0 | 0 |
| **Rolling-circles** | 105448 | 13627326 | 0.62 |
| **Unclassified:** | 1156 | 181522 | 0.01 |
| **Total interspersed repeats:** | na | 540592282 | 24.78 |
| **Small RNA:** | 10052 | 995117 | 0.05 |
| **Satellites:** | 3639 | 1433924 | 0.07 |
| **Simple repeats:** | 616339 | 36363820 | 1.67 |
| **Low complexity:** | 71663 | 7022592 | 0.32 |

**Supplementary Table 13. Content of repeat elements in *Anolis auratus* genome.**

| | Number of elements | Length occupied (bp) | Percentage of sequence |
|---|---|---|---|
| **Retroelements** | 1916123 | 461310730 | 22.86 |
| **SINEs:** | 348524 | 54403272 | 2.7 |
| **Penelope** | 221123 | 39878904 | 1.98 |
| **LINEs:** | 1472571 | 369879139 | 18.33 |
| **CRE/SLACS** | 0 | 0 | 0 |
| **L2/CR1/Rex** | 856048 | 208121218 | 10.32 |
| **R1/LOA/Jockey** | 437 | 107772 | 0.01 |
| **R2/R4/NeSL** | 33915 | 10099122 | 0.5 |
| **RTE/Bov-B** | 215741 | 57679462 | 2.86 |
| **L1/CIN4** | 87788 | 35473264 | 1.76 |
| **LTR elements:** | 95028 | 37028319 | 1.84 |
| **BEL/Pao** | 23666 | 7284004 | 0.36 |
| **Ty1/Copia** | 7757 | 3421414 | 0.17 |
| **Gypsy/DIRS1** | 43706 | 18691097 | 0.93 |
| **Retroviral** | 8676 | 2617411 | 0.13 |
| **DNA transposons** | 970996 | 152701852 | 7.57 |
| **hobo-Activator** | 506939 | 84786861 | 4.2 |
| **Tc1-IS630-Pogo** | 297708 | 46646253 | 2.31 |
| **En-Spm** | 0 | 0 | 0 |
| **MuDR-IS905** | 0 | 0 | 0 |
| **PiggyBac** | 248 | 57277 | 0 |
| **Tourist/Harbinger** | 112533 | 14584867 | 0.72 |
| **Other** | 0 | 0 | 0 |
| **Rolling-circles** | 66825 | 8485753 | 0.42 |
| **Unclassified:** | 904 | 140358 | 0.01 |
| **Total interspersed repeats:** | na | 614152940 | 30.44 |
| **Small RNA:** | 12891 | 1083587 | 0.05 |
| **Satellites:** | 2573 | 189000 | 0.01 |
| **Simple repeats:** | 882277 | 120889949 | 5.99 |
| **Low complexity:** | 70234 | 6324004 | 0.31 |

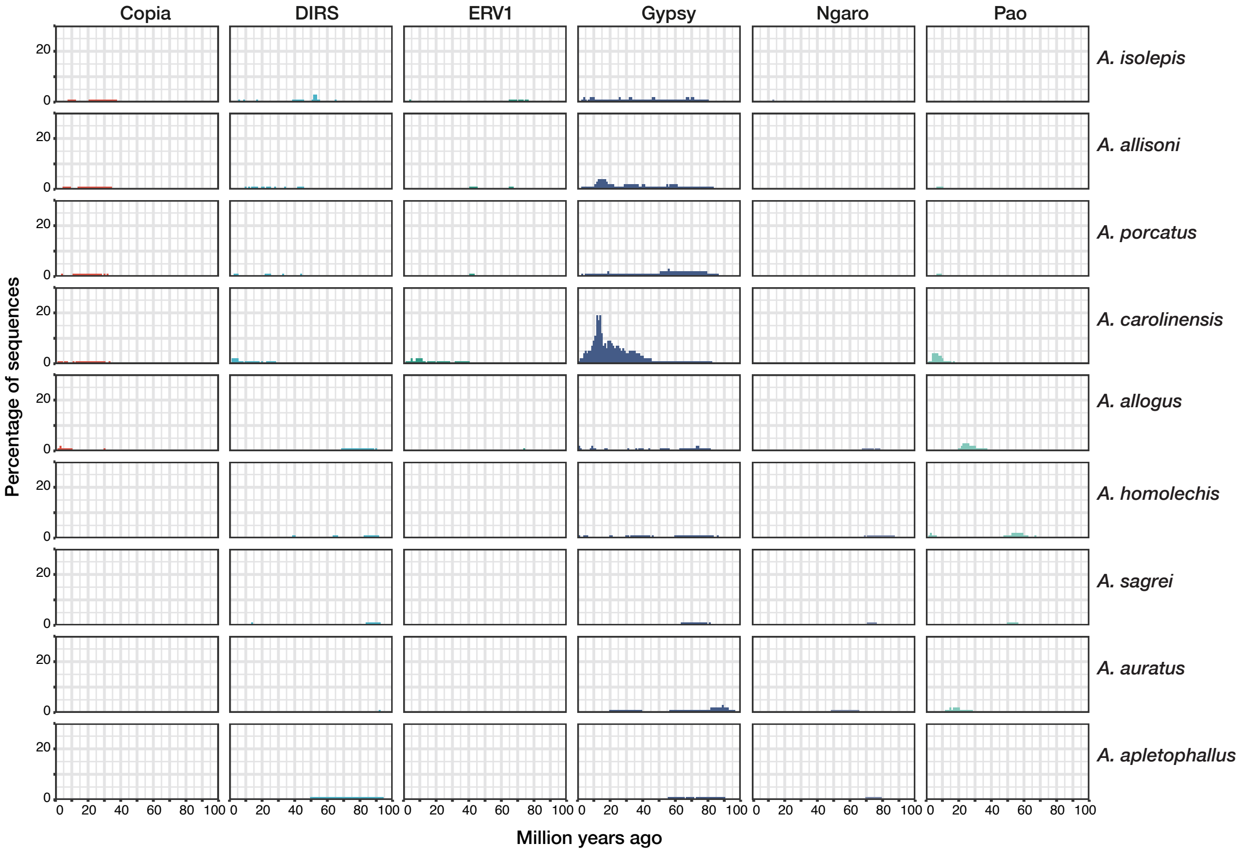


**Supplementary Fig. 1. Repeat landscape for each LTR transposon family of *Anolis* lizards included in this study.**

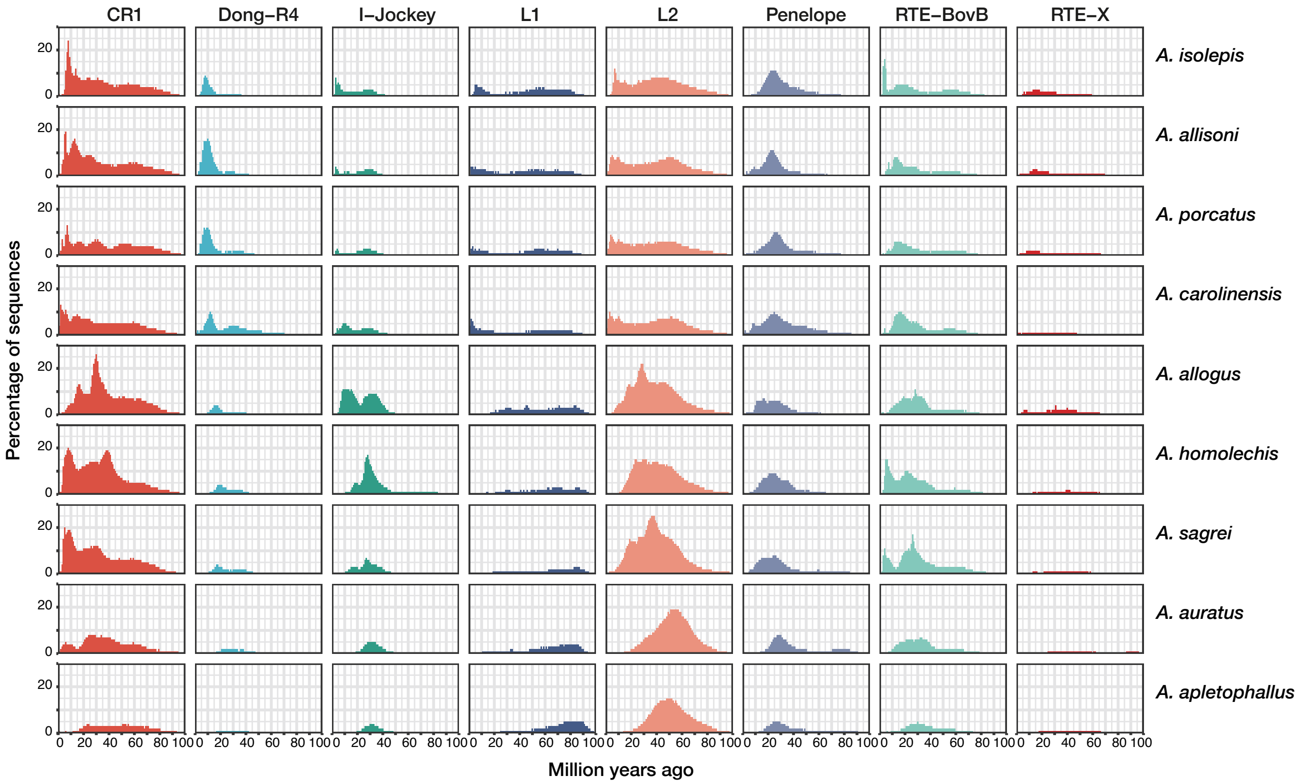


**Supplementary Fig. 2. Repeat landscape for each LINE transposon family of *Anolis* lizards included in this study.**

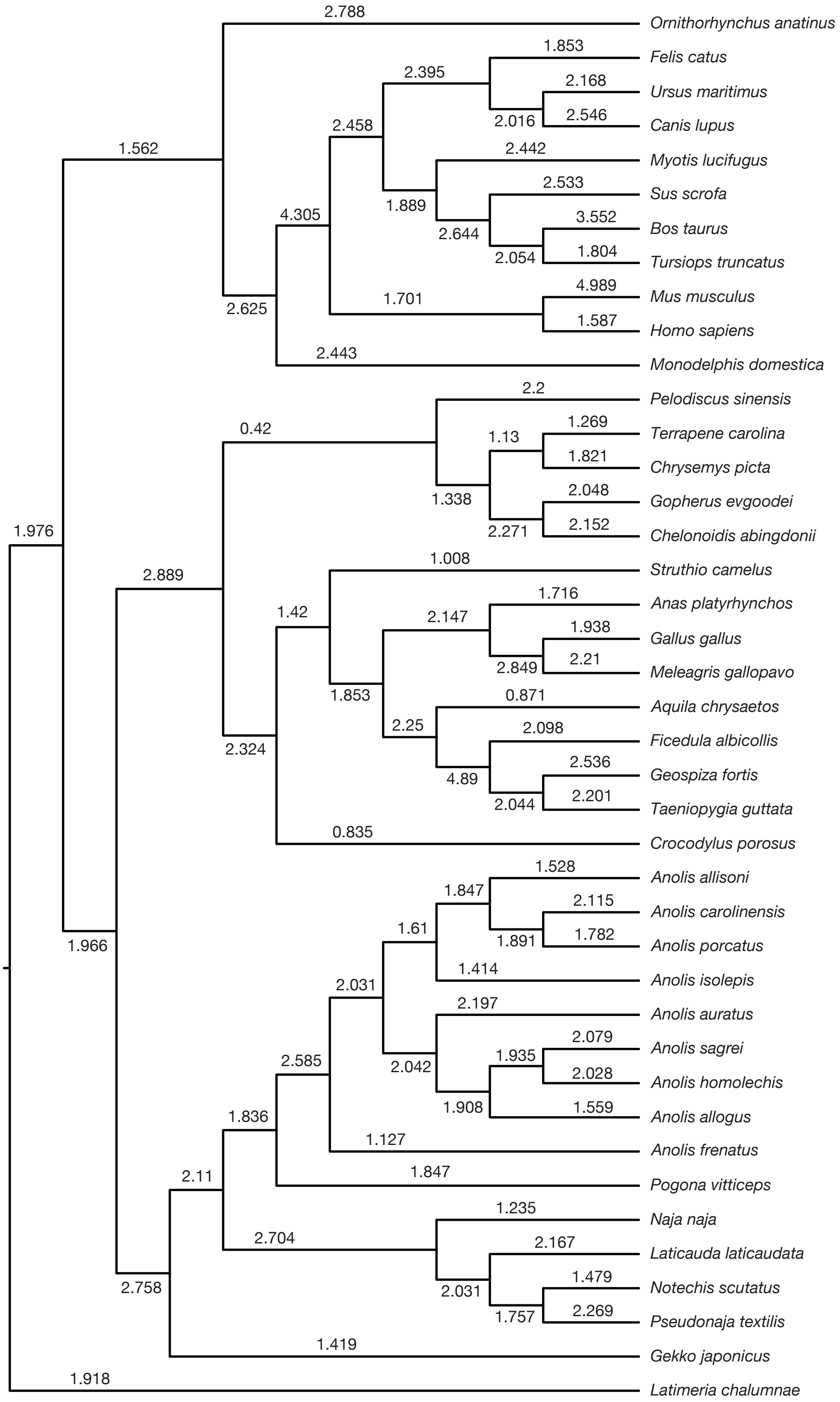

Ornithorhynchus anatinus
Felis catus
Ursus maritimus
Canis lupus
Myotis lucifugus
Sus scrofa
Bos taurus
Tursiops truncatus
Mus musculus
Homo sapiens
Monodelphis domestica
Pelodiscus sinensis
Terrapene carolina
Chrysemys picta
Gopherus evgoodei
Chelonoidis abingdonii
Struthio camelus
Anas platyrhynchos
Gallus gallus
Meleagris gallopavo
Aquila chrysaetos
Ficedula albicollis
Geospiza fortis
Taeniopygia guttata
Crocodylus porosus
Anolis allisoni
Anolis carolinensis
Anolis porcatus
Anolis isolepis
Anolis auratus
Anolis sagrei
Anolis homolechis
Anolis allogus
Anolis frenatus
Pogona vitticeps
Naja naja
Laticauda laticaudata
Notechis scutatus
Pseudonaja textilis
Gekko japonicus
Latimeria chalumnae
2.788
1.853
2.395
2.168
2.016
2.546
2.458
2.442
1.562
1.889
2.533
4.305
2.644
3.552
2.054
1.804
1.701
4.989
2.625
1.587
2.443
2.2
0.42
1.269
1.13
1.821
1.338
2.048
1.976
2.271
2.152
1.008
2.889
1.716
1.42
2.147
1.938
2.849
2.21
1.853
0.871
2.25
2.098
2.324
2.536
4.89
2.044
2.201
0.835
1.528
1.847
2.115
1.61
1.891
1.782
1.966
1.414
2.031
2.197
2.079
2.042
1.935
2.028
2.585
1.908
1.559
1.836
1.127
2.11
1.847
1.235
2.704
2.167
1.479
2.758
2.031
1.757
2.269
1.419
1.918

**Supplementary Fig. 3. DNA substitution rate (substitutions per billion years) for each branch of phylogenetic tree of sarcopterygian vertebrates.**

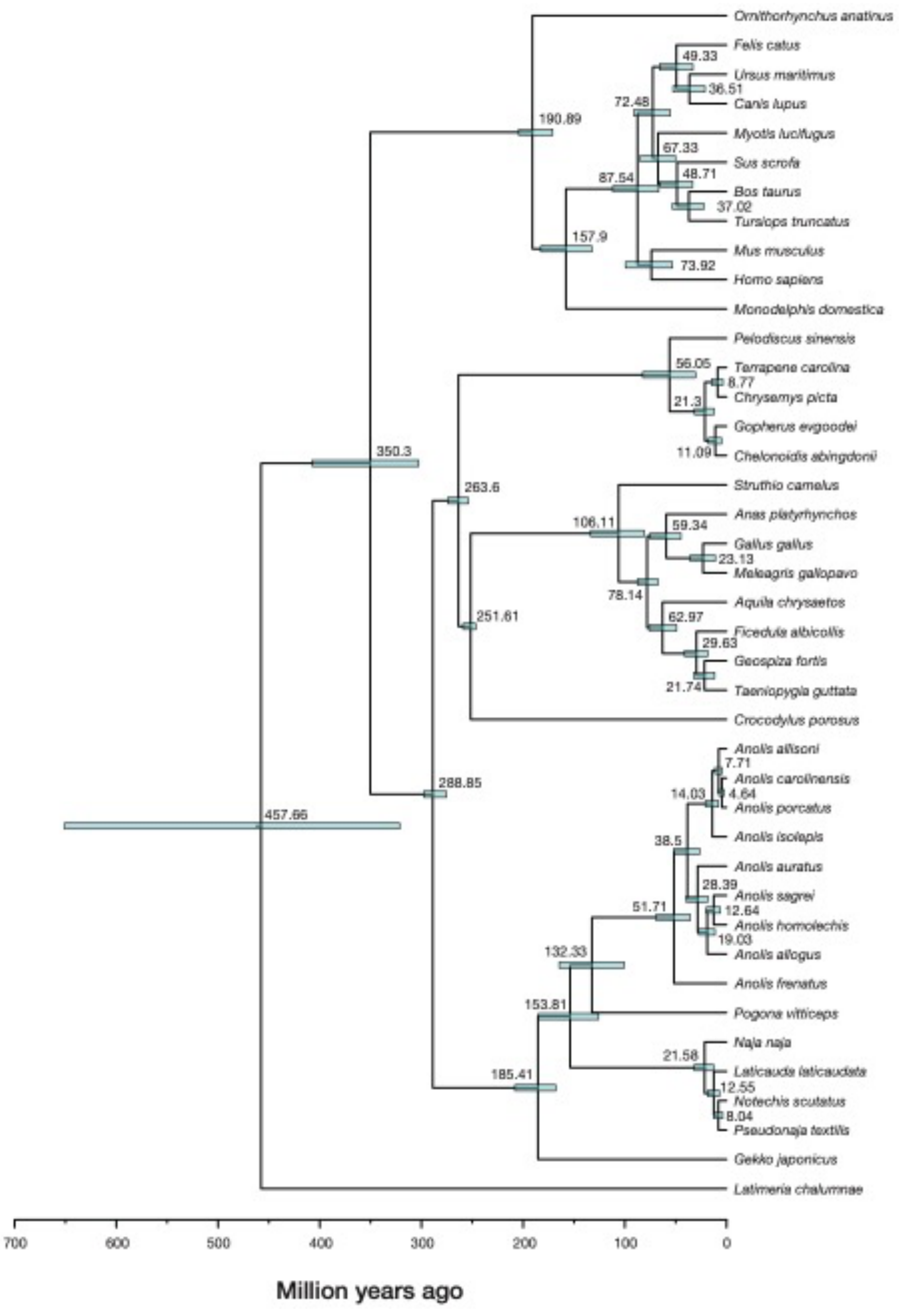

Ornithorhynchus anatinus
Felis catus
Ursus maritimus
Canis lupus
Myotis lucifugus
Sus scrofa
Bos taurus
Tursiops truncatus
Mus musculus
Homo sapiens
Monodelphis domestica
Pelodiscus sinensis
Terrapene carolina
Chrysemys picta
Gopherus evgoodei
Chelonoidis abingdonii
Struthio camelus
Anas platyrhynchos
Gallus gallus
Meleagris gallopavo
Aquila chrysaetos
Ficedula albicollis
Geospiza fortis
Taeniopygia guttata
Crocodylus porosus
Anolis allisoni
Anolis carolinensis
Anolis porcatus
Anolis isolepis
Anolis auratus
Anolis sagrei
Anolis homolechis
Anolis allogus
Anolis frenatus
Pogona vitticeps
Naja naja
Laticauda laticaudata
Notechis scutatus
Pseudonaja textilis
Gekko japonicus
Latimeria chalumnae
49.33
36.51
72.48
67.33
48.71
37.02
87.54
157.9
73.92
190.89
56.05
8.77
21.3
11.09
350.3
263.6
106.11
59.34
23.13
78.14
251.61
62.97
29.63
21.74
7.71
4.64
14.03
288.85
457.66
38.5
28.39
12.64
19.03
51.71
132.33
153.81
21.58
12.55
8.04
185.41
700
600
500
400
300
200
100
0
Million years ago

**Supplementary Fig. 4 Divergence time tree of sarcopterygian vertebrates reconstructed by Bayesian method implemented in MCMCTree.** Node bars indicate 95% highest posterior density.

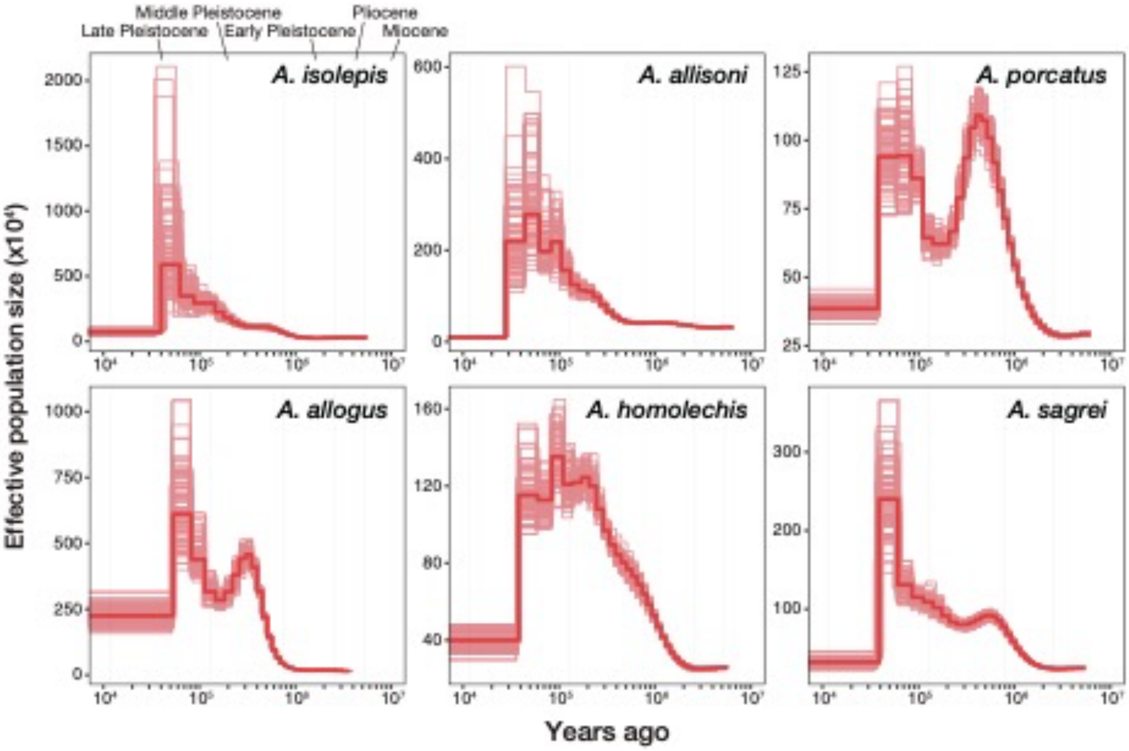


**Supplementary Fig. 5. Past effective population size of six Cuban *Anolis* lizards inferred using PSMC with bootstrap results.** The thick red lines are the consensus result, and the thin light red lines are each bootstrap result.
